# Spatiotemporal gene expression atlas of the extremophyte *Schrenkiella parvula*

**DOI:** 10.1101/2022.10.24.513627

**Authors:** Chathura Wijesinghege, Guannan Wang, Pramod Pantha, Kieu-Nga Tran, Maheshi Dassanayake

## Abstract

Extremophytes are naturally selected to survive environmental stresses, but scarcity of genetic resources for them developed with spatiotemporal resolution limit their use in stress biology. *Schrenkiella parvula* is one of the leading extremophyte models with initial molecular genomic resources developed to study its tolerance mechanisms to high salinity. Here we present a transcriptome atlas for *S. parvula* with subsequent analyses to highlight its diverse gene expression networks associated with salt responses. We included spatiotemporal expression profiles, expression specificity of each gene, and co-expression and functional gene networks representing 115 transcriptomes sequenced from 35 tissue and developmental stages examining their responses before and after 27 salt treatments in our current study. The highest number of tissue-preferentially expressed genes were found in seeds and siliques while genes in seedlings showed the broadest expression profiles among developmental stages. Seedlings had the highest magnitude of overall transcriptomic responses to salinity compared to mature tissues and developmental stages. Differentially expressed genes in response to salt were largely mutually exclusive but shared common stress response pathways spanning across tissues and developmental stages. Our foundational dataset created for *S. parvula* representing a stress-adapted wild plant lays the groundwork for future functional, comparative, and evolutionary studies using extremophytes aiming to uncover novel stress tolerant mechanisms.

**SIGNIFICANCE STATEMENT:** Concerted transcriptomic responses coordinated across developmental stages and tissues are required to complete a plant lifecycle under salt stress. Transcriptomic resources created with spatiotemporal resolution for plants are rare and for stress-adapted plants rarer. We present a transcriptome atlas enabling discovery of genes and networks evolved as adaptations to salt stress in a model extremophyte, *Schrenkiella parvula*. The spatiotemporally resolved gene expression networks are largely non-overlapping but functionally connected through synergistic stress responsive pathways.

## INTRODUCTION

Salt stress is a complex stress and plants have evolved diverse responses to salt stress (Munns *et al*., 2016; Flowers *et al*., 2010; Pantha and Dassanayake, 2020). Less than five percent of land plants are capable of resilient growth under high salinities represented by extremophytes (Santos *et al*., 2016). Extremophytes are broadly categorized as plants highly tolerant of multiple environmental stresses (Barak and Farrant, 2016). Increasing soil salinities are always growth inhibitory to plants at some intensity but extremophytes show a higher tolerance capacity (Santiago-Rosario *et al*., 2021; Zhang *et al*., 2018). Therefore, extremophyte relatives of crops not just provide insight on genetic variation that is frequently lost through domestication processes, but are also excellent systems to study genetic mechanisms evolved to survive harsh environments (Eshel *et al*., 2022; Kumar *et al*., 2021; Birkeland *et al*., 2020; Amin *et al*., 2019). However, only a handful of extremophytes have been developed as molecular genetic models to systematically study stress tolerance. Their use in plant stress biology is increasing with our increasing need to design resilient crops to survive environmental stress during a climate crisis (Cushman *et al*., 2022).

*Schrenkiella parvula* (Schrenk) German & Al-Shehbaz (former names: *Thellungiella parvula* and *Eutrema parvulum*) is a leading extremophyte model in the Brassicaceae family closely related to mustard crops (Zhu, 2015; Kazachkova *et al*., 2018; Oh *et al*., 2012). It is highly tolerant of sodium and other salts that contribute to soil salinity (Oh *et al*., 2014). *S. parvula* is naturally found in the Irano-Turanian region often near saline lakes (Hajiboland *et al*., 2018; Tug *et al*., 2008). Its recent development as a functional genomic model has resulted in a high quality chromosome-level genome (Dassanayake *et al*., 2011), transformation methods (Wang *et al*., 2019), physiological assessments (Orsini *et al*., 2010; Tran *et al*., 2021), and multi-omics datasets generated to investigate adaptations to environmental stress (Tran *et al*., 2022; Pantha *et al*., 2021; Wang *et al*., 2021; Wijesinghege *et al*., 2022; Oh *et al*., 2014). Prior studies have also focused on landmark genes in salt stress regulation but with novel adaptations that make their function different in *S. parvula* to those in salt-sensitive plants (Ali *et al*., 2016; Ali *et al*., 2018; Jarvis *et al*., 2014). Multiple comparative studies between *S. parvula* and *Arabidopsis thaliana* have shown that these plants have highly diverged gene expression responses even for well-known stress regulatory networks when treated with equivalent salt levels (Sun *et al*., 2022; Tran *et al*., 2022; Oh *et al*., 2014). Therefore, *S. parvula* has been proposed as an excellent wild crop relative to mine genetic variation used for crop development in marginal lands due to natural salinization, climate change driven land salinization, or irrigation malpractices (Zhang *et al*., 2018; Himabindu *et al*., 2016; Isayenkov, 2019; Rawat *et al*., 2022).

The majority of genetic experiments exploring salt tolerance have used short term salt stress given to seedlings, bulk roots, or bulk shoots at vegetative growth phases and also have used model plants or crops not recognized for their salt tolerance (Munns, 2005; Negrão *et al*., 2017; Zhu *et al*., 1998). However, transcriptomic responses coordinated across developmental stages and tissues leading to physiological studies show that plants respond to salt with tissue and developmental stage specificity (Negrão *et al*., 2017). The corresponding input from gene expression studies to investigate salt tolerance at a spatiotemporal scale exists for the model plant *A. thaliana* (Cheng *et al*., 2017), but not for any plant capable of salt resilient growth. Therefore, a transcriptome atlas generated for an extremophyte model, *S. parvula* provides an exemplary resource to fill a critical gap in our knowledgebase to study tissue and developmental salt tolerance at the whole plant scale.

Here we provide a transcriptome atlas for *S. parvula* with over thirty tissues and developmental stages profiled to highlight their transcriptome responses in salt treated compared to control conditions. Our results indicate that *S. parvula* transcriptomes are highly spatiotemporally distinct with non-overlapping differently co-expressed gene modules. However, the mutually exclusive salt responsive gene modules spanning developmental stages and tissues highly overlap in functional pathways representative of stress responses.

## RESULTS

### Spatiotemporally distinct salt responsive transcriptomes of *Schrenkiella parvula*

To characterize the spatiotemporal gene expression profiles and salt stress responses among tissues and developmental stages in the extremophyte *Schrenkiella parvula*, we sequenced RNA from tissues, organs, or entire plants at different developmental stages. Our sampling scheme was designed to capture all major developmental stages and how each phase responded to high salinity during *S. parvula*’s lifecycle from seeds, seedlings, mature plants before flowering, to reproductive stages (including flowers and siliques that developed before and after salt treatments) (Figure 1 and S1). Detailed descriptions on sample and salt treatment conditions are given in Table S1 and Experimental procedures.

**Figure 1.**
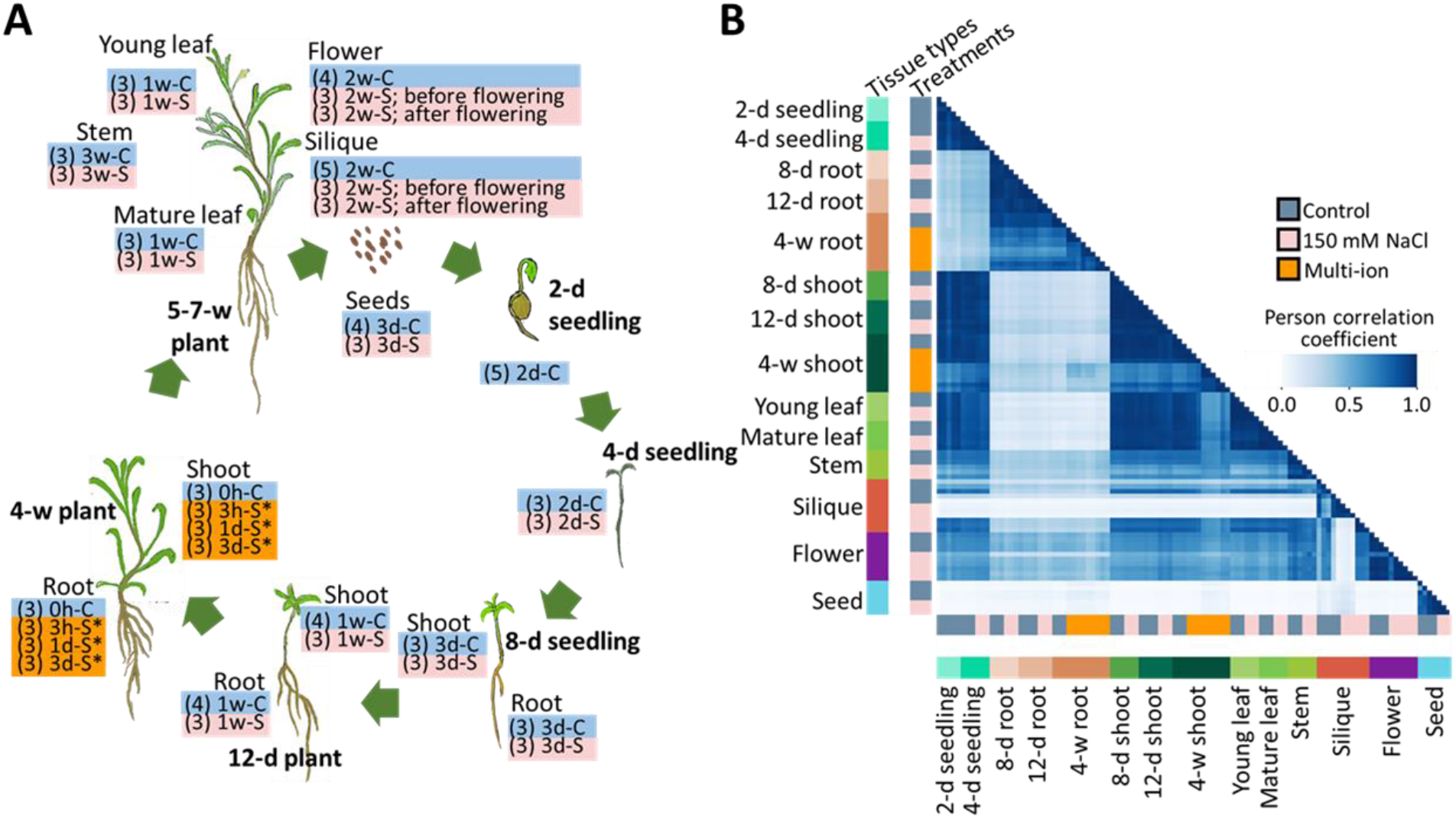
Generation and overview of the *Schrenkiella parvula* transcriptome atlas. **[A]** Transcriptome atlas created from RNA samples extracted throughout the lifecycle. Age of the plant when samples were taken is given at each developmental stage. Organ/tissue type is given for each set of samples that contain control or salt treatments outlined within a box. Control (C) condition did not have added NaCl. Two types of salt treatments were given: (a) 150 mM NaCl (S) and (b) multi-ion salt (S*) given as a combined stress consisting of 250 mM NaCl, 250 mM KCl, 30 mM LiCl, and 15 mM H_3_BO_3_. Duration of treatments ranged from 3 hours to 3 weeks. h-hour, d-day, and w-week. Stems harvested from 4^th^-6^th^ internodes. Young leaves consisted of leaf primodia and the youngest detectable leaf. Mature leaves were 5^th^ to 7^th^ from the base. Two sets of flowers and siliques were harvested with salt treatments started before and after flowering. Seeds were imbibed in control or salt solutions. Biological replicates (3-5) used per condition is given in parenthesis within boxes to show treatment and duration. **[B]** Pearson correlation calculated based on the expression of genes between each of the samples and their replicates. Colored bars on x- and y- axes indicate the sample types and treatments.

We generated 107 RNAseq samples representative of 33 distinct conditions (Figure 1) with an average of 69 million reads per sample (Table S1). We assessed the overall transcriptome similarity across samples using Pearson correlation calculated for gene expression between all sample pairs (Figure 1B). Biological replicates showed high similarity within each sample group while samples from major organ types clustered together (Figure S2). Shoots, roots, and reproductive tissues (flowers, siliques, and seeds) formed distinct transcriptome clusters. Gene expression distributions were created only with those genes that had expression values ≥ 2 TPM (transcripts per million) in at least one sample to avoid low-expressed genes with high variance that may add to responses poorly supported at the overall transcriptome level. Only protein coding genes in the *S. parvula* reference genome v2.2 (Phytozome genome ID: 574) were quantified in our analyses and will be referred to as genes hereafter.

Each tissue-developmental sample consisted of 12,000 to 17,000 (43 - 67% in the genome) expressed genes (Figure 2A, Table S2). Seeds showed the lowest total number of expressed genes among samples (Figure 2A), but its overall transcriptome expression distribution was comparable to other tissues (Figure 2B). Most tissues and developmental stages had over 60% of genes expressed regardless of control or salt treated conditions and 40% of those were expressed in all tissues (12 control and 13 salt sample types; Figure 2C).

**Figure 2.**
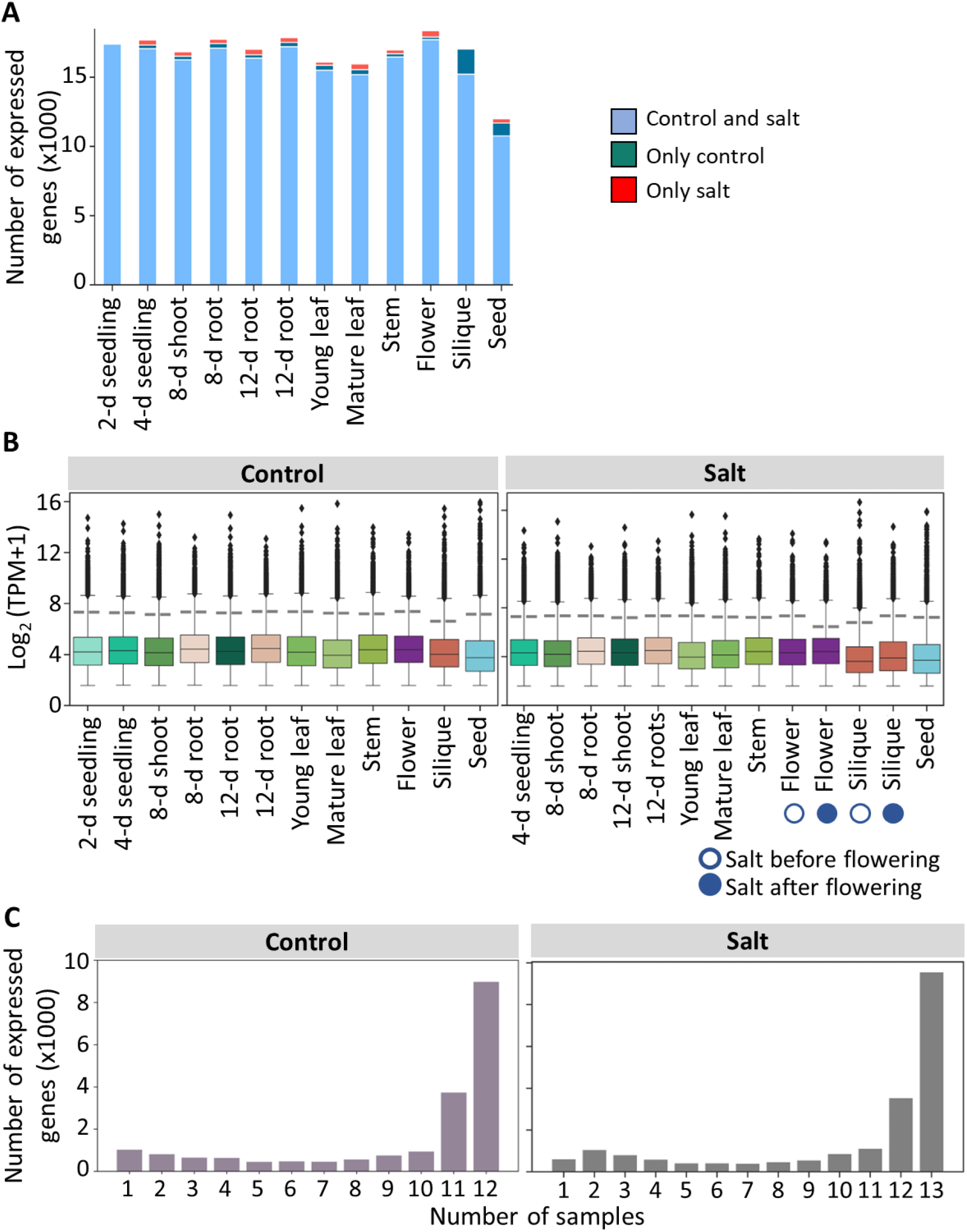
Spatiotemporal gene expression distribution in *S. parvula*. **[A]** Number of genes expressed in each sample. An expressed gene per sample was counted if the average expression within that sample was ≥ 2 TPM. **[B]** Expression distribution of genes in different tissues and developmental stages under control (left) and salt treated (right) conditions. **[C]** Number of expressed genes that are shared among samples under (left) and salt treated (right) conditions.

We developed the entire transcriptome atlas dataset into a searchable expression browser as a resource for plant stress biologists accessible at https://www.lsugenomics.org/resources. In this portal users can interactively explore individual gene expression profiles quantified spatiotemporally. A screenshot of the user interface is shown in Figure S3. RNAseq data and counts from our study can be accessed via the US Department of Energy Joint Genome Institute Genome portal accessible at https://genome.jgi.doe.gov/portal/SchparEProfiling/SchparEProfiling.info.html under Project ID: 1263770 with other genome visualization tools available through Phytozome.

### Distinct gene co-expression networks reflect major developmental processes and functions associated with tissues

We built gene co-expression networks using WGCNA (Langfelder and Horvath, 2008) to identify distinct gene expression modules that span across tissues and developmental stages in *S. parvula* (Figure 3). We included only control samples and also excluded bulk roots and shoots from 4-week old plants (that contained pooled leaves of all ages and stems) as the input dataset to generate the co-expressed networks. We identified 21 non-overlapping co-expression gene modules (Figure 3A). All gene modules with their spatiotemporal expression distribution and enriched functions are given in (Figure 3B and S4). The largest module (Module-1) consisted of 7,693 genes which were enriched in organelle organization and carbohydrate metabolism (Figure S4). The second largest (Module-2) was preferentially expressed in green tissues and was enriched in processes associated with photosynthesis. The smaller modules showed increased specificity in their expression profiles. For example, Module-14 was preferentially expressed in roots, 2- and 4-day-old seedlings with enriched functions involved in root morphogenesis, while Module-16 was preferentially expressed in siliques and seeds with functions representative of ABA responses and seed maturation (Figure 3B). Overall, the *S. parvula* gene co-expression network showed coordinated developmental processes across tissue types characteristic of the functions those tissues are expected to perform.

**Figure 3.**
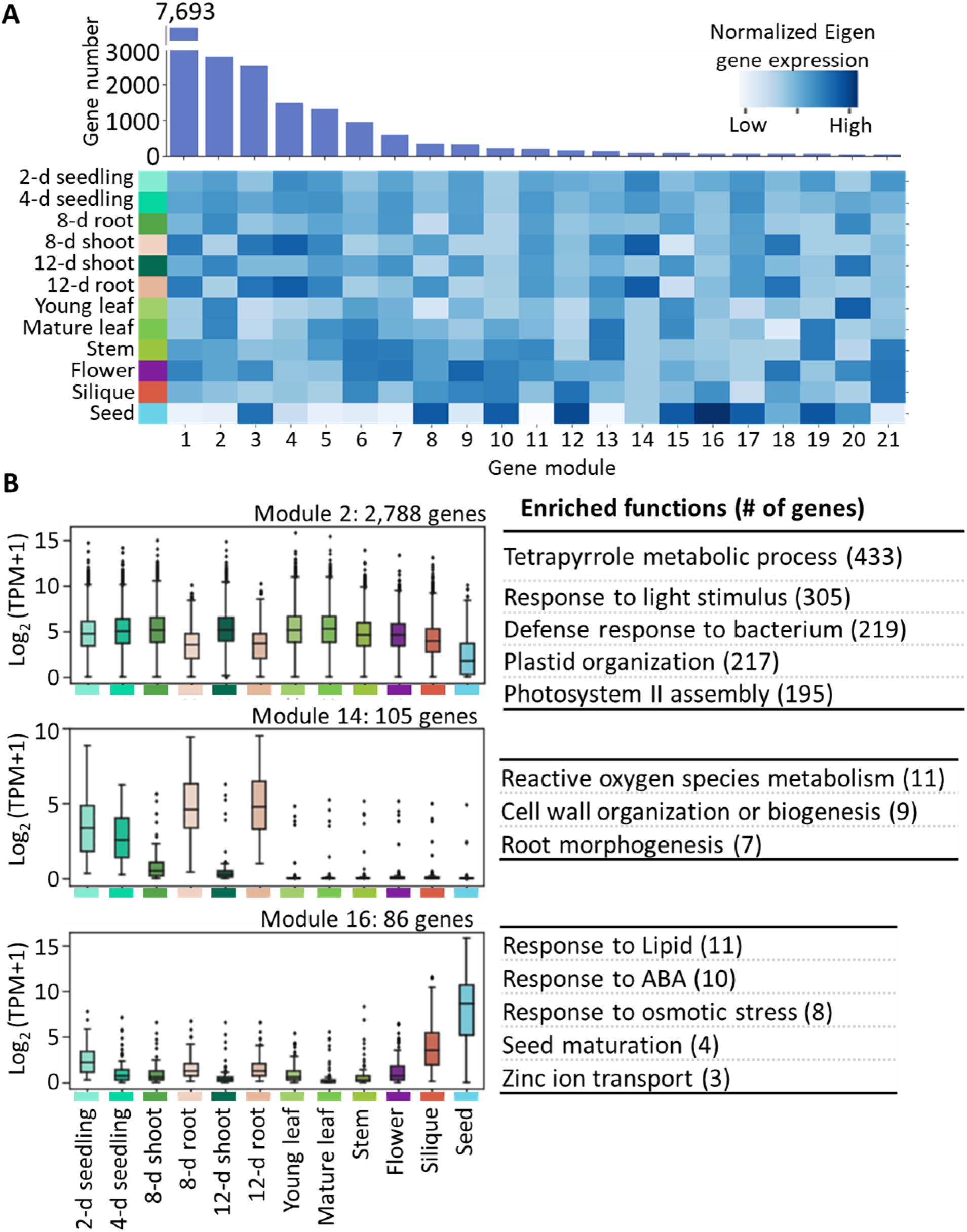
Gene co-expression network across tissues and developmental stages. **[A]** Co-expression gene network shown in 21 distinct gene modules sorted by their size from largest to smallest. The upper panel shows the number of genes per module. The lower panel shows the normalized expression of a representative gene per module (Eigen genes) and how each module is spatiotemporally expressed. **[B]** Expression distribution of selected co-expression modules. Module 2 is an example of a gene group co-expressed in all samples while Modules 14 and 16 show expression mainly in few samples. Functional processes enriched in each module are shown on the right.

### Preferential expression of genes in tissues and developmental stages

Genes with preferential expression in tissues or developmental stages give rise to the specificity of spatiotemporal expression of the genome. Highly expressed genes with preferential expression in limited tissues or developmental stages can serve as marker genes for distinct stress responses specific to the condition. Therefore, we calculated the Tissue Specificity Index (τ) (Yanai *et al*., 2005) independently for control and salt treated samples. Bulk roots and shoots from 4-week old plants were excluded to allow the identification of more specific expression assigned to genes expressed in the rest of the samples. Specific gene expression measured as τ ranges from 0 to 1, where 0 indicates that a gene is broadly expressed in all conditions, while 1 indicates its expression is restricted to one condition. We considered τ ≥ 0.8 to represent highly tissue- or developmental stage-preferentially expressed genes in our study (comparable to tissue-specific genes identified for human tissues (Yanai *et al*., 2005)). We refer to these as preferentially expressed genes (PEGs).

We observed that reproductive tissues (flowers, siliques, and seeds) had a higher number of PEGs, while young leaves and seedlings had a lower number of PEGs among all samples (Figure 4A). It is interesting to note that 8-day shoots had far fewer PEGs compared to the 8-day root tissue derived from the same seedling pool. Not all PEGs had high expression and could potentially serve as marker genes for tissues or developmental stages in *S. parvula* (Figure 4B). Therefore, we selected the most highly expressed four PEGs per condition based on their relative expression within each sample as a candidate set of tissue and developmental stage marker genes for *S. parvula* (Figure S5).

**Figure 4.**
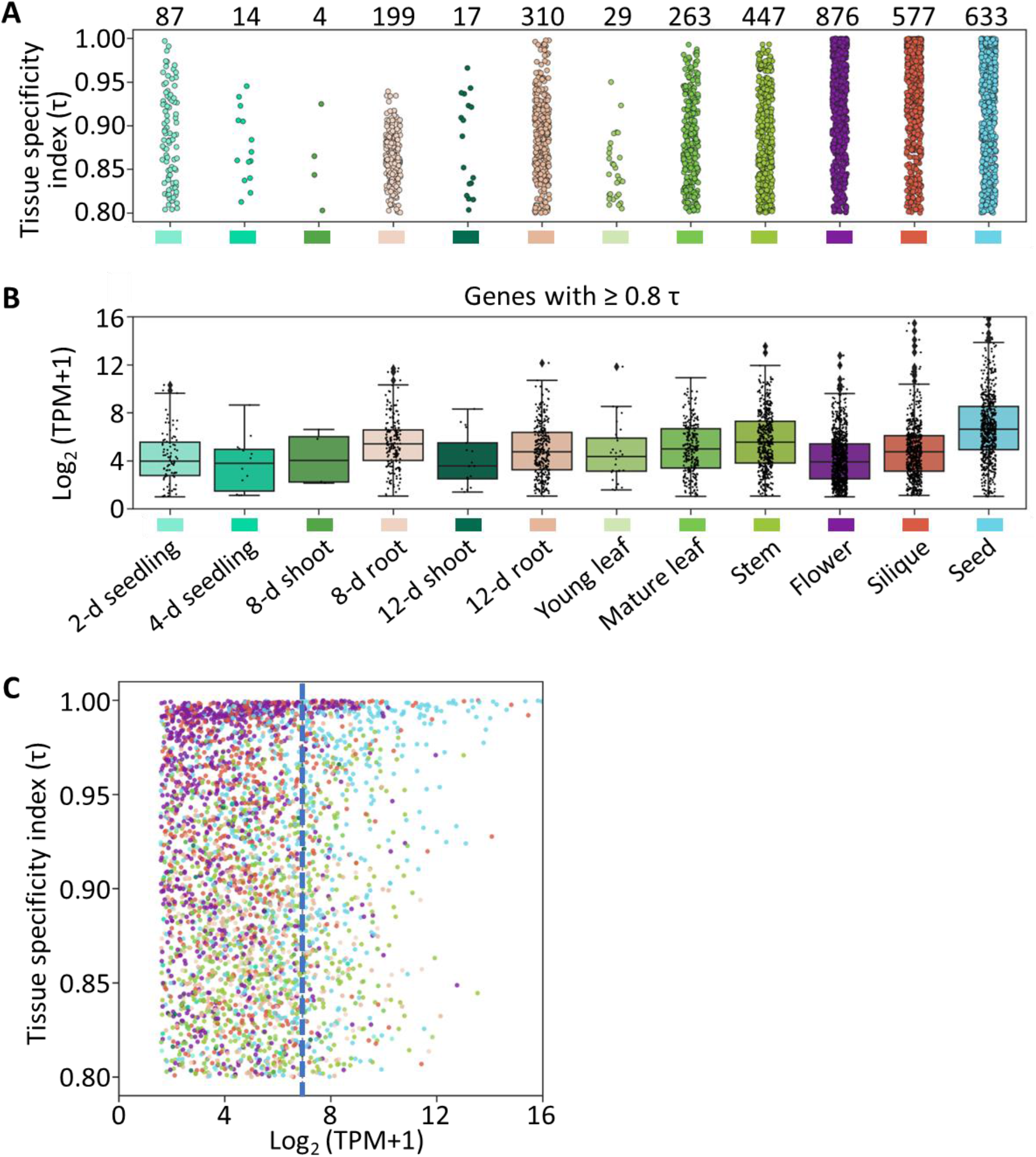
Preferential expression of genes in tissues or developmental stages in control samples. **[A]** Genes with tissue specificity index (τ) ≥ 0.8. **[B]** Expression distribution of preferentially expressed genes among samples. **[C]** Highly tissue or developmentally restricted expression and expression strength of genes. Dots are colored corresponding sample colors given in A and B. Dashed line indicates the expression cutoff for top 5% of highly expressed genes with high preferential expression specificity.

We did not find a strong correlation between preferential expression and expression strength of a gene (R^2^ = 0.0081 at *p* ≤ 0.01) (Figure 4C). However, ∼46% of PEGs that had the highest expression (top 5% expressed) were found in seeds (Figure 4C). PEGs from stems (19% of PEGs) and siliques (10% of PEGs) were also among the top 5% highly expressed PEGs (Table S3).

It is generally accepted that transcription factors are important in regulating the spatiotemporal gene expression and maintaining tissue identity. Therefore, we tested if transcription factors had higher τ values than other genes. Indeed, transcription factor genes were significantly more preferentially expressed than other genes (Figure 5A). Next, we examined if transcription factors were enriched among PEGs in each sample (Figure 5B). Many of the tissues and developmental stages examined were enriched in transcription factors among PEGs (marked by an asterisk in the upper plot of Figure 5B). Expression strength of transcription factor genes in all tissues, however, were generally lower than the other genes (lower plot of Figure 5B). We did not detect transcription factors among PEGs for 4-day seedlings and shoots of 8-day seedlings. The expression profile for each transcription factor identified as a PEG showed that they were expressed with minimal overlaps in adjacent tissues or development stages except in 8-day and 12-day old roots (Figure 5C).

**Figure 5.**
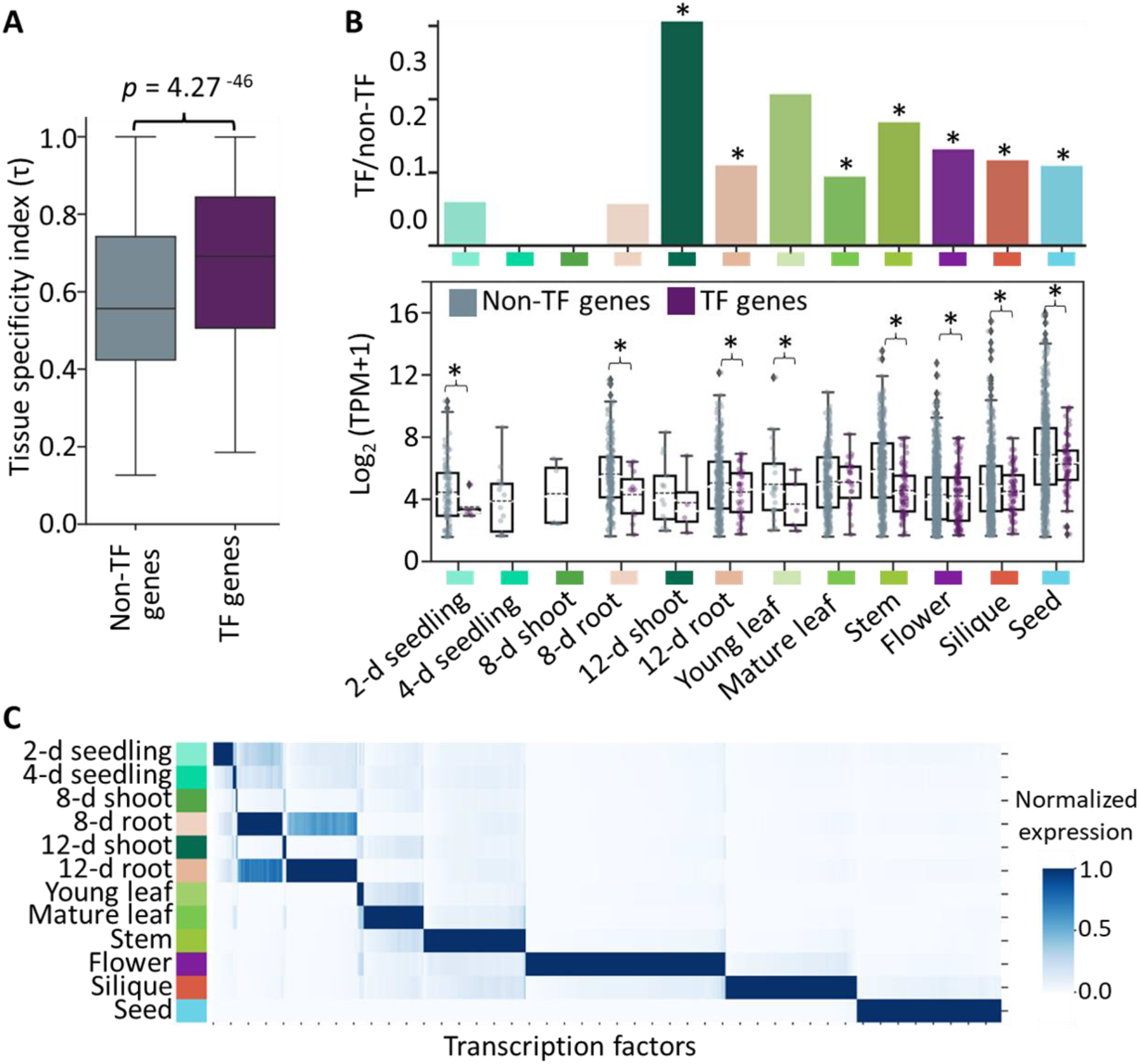
Spatiotemporal expression of genes coding for transcription factors. **[A]** Expression specificity of transcription factors (TF) and non-transcription factors (Non-TF) measured by tissue specificity index (τ). *p*-value based on Wilcoxon test applied to two groups. **[B]** Enrichment and expression strength of TF genes when compared to non-TF genes in control samples. All genes considered had τ ≥ 0.8. Upper plot: the number of TF genes as a ratio between TF and non-TF genes expressed in each sample. Asterisks indicate significant differences determined by a test for over-representation using a hypergeometric *p*-value ≤ 0.05. Lower plot: Expression distribution of TFs and Non-TFs across samples. Asterisks indicate significant difference determined by Wilcox test at *p* ≤ 0.05. **[C]** Normalized expression of TFs preferentially expressed (τ ≥ 0.8) in tissues and developmental stages.

### Gene expression specificity shifts under high salinity

Given that salt stress leads to a systemic response across tissues and developmental stages in *S. parvula* (Tran *et al*., 2021), we wanted to identify genes that responded to salt by changing their expression profile specificity using shifts in τ. A large proportion of *S. parvula* genes did not shift their expression specificity when transitioning from control to salt treated conditions (Figure 6A). We categorized the genes that responded to salt by switching tissue specificity into two groups: (1) tissue specificity lost - 125 genes switched from high to low τ upon salt treatment and (2) tissue specificity gained - 129 genes switched from low to high τ upon salt treatment.

**Figure 6.**
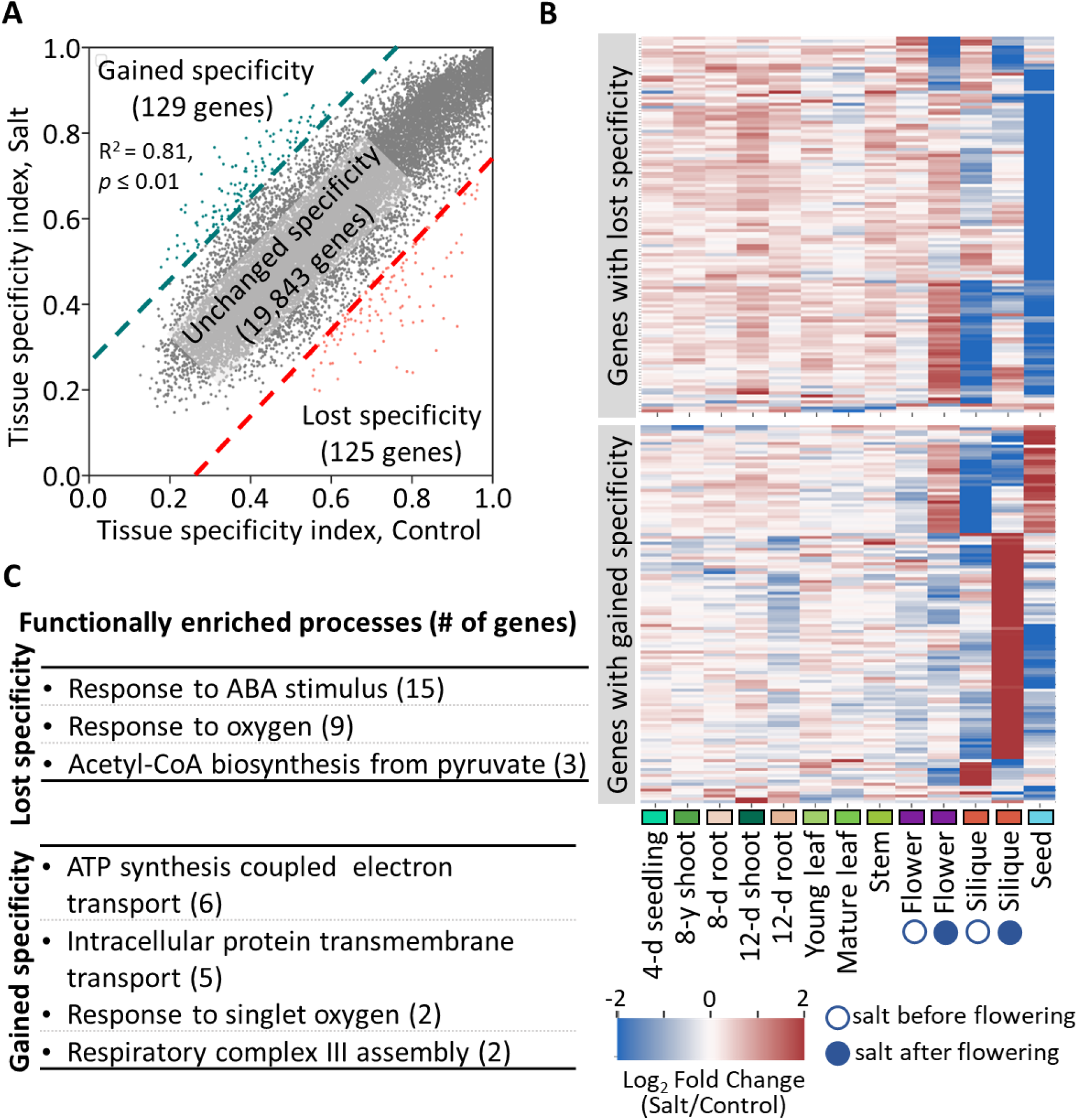
Gene expression specificity switches when transitioning from control to salt. **[A]** Tissue specificity index (τ) per gene under control and salt treated samples. Dash lines indicate the difference between τ_control_ and τ_salt_ ≥ 0.25. Green points indicate genes that increased their expression specificity and red points indicate genes that decreased their expression specificity under salt. **[B]** The expression change between control and salt as a fold change for genes highlighted in green and red given in panel A. **[C]** Functional processes enriched among genes that changed their expression specificity under salt (genes considered for panel B).

Notably, the genes that had high τ values, but decreased them under high salinity were predominantly expressed (80%) in seeds. Moreover, these genes in seeds had higher expression under control conditions but were downregulated in response to salt (upper panel in Figure 6B). Contrastingly, the same genes when expressed in other samples had lower expression in control conditions, but were induced under high salinity (Figure 6B). This group of genes that lowered their tissue specificity under high salinity were enriched for ABA stress responsive genes (Figure 6C). Thus, our results suggest that upon exposure to salt, stress responsive genes regulated under ABA are broadly expressed from their more restricted expression in seeds and are also overall induced in most tissues while being suppressed in seeds.

Plants treated with salt before the onset of flowering and those that were treated with salt after the onset of flowering had very distinct transcriptome responses for flowers and siliques despite the similar tissue types used to obtain the transcriptomes (Figure 6B). The genes that increased in τ upon salt were mostly expressed in siliques developed in plants that were salt treated after the onset of flowering (Figure 6B). The gain in tissue specificity upon high salinity was mainly due to induced gene expression in flowers and siliques that were treated with salt after flowering and genes induced in seeds. It should be noted that these genes between siliques and seeds do not overlap and show contrasting paths to gaining expression specificity under high salinity (Figure 6B).

### Duplicated genes show higher expression specificity than single copy genes

Duplicated genes are known to show more specialized tissue specific functions than single-copy genes in multicellular organisms (Freilich *et al*., 2006). Therefore, we predicted that multi-copy genes will likely show more tissue specific expression. We found that multi-copy genes in *S. parvula* are expressed at higher τ values than found with single-copy genes (Figure 7A). Next, we wanted to identify duplicated genes which had both copies categorized as PEGs (τ ≥ 0.8) from those duplicated genes that had neither copy showing preferential expression in tissues (Figure 7B). The more specialized duplicated genes are enriched in functions associated with lipid metabolism and organ development (Figure 7C).

**Figure 7.**
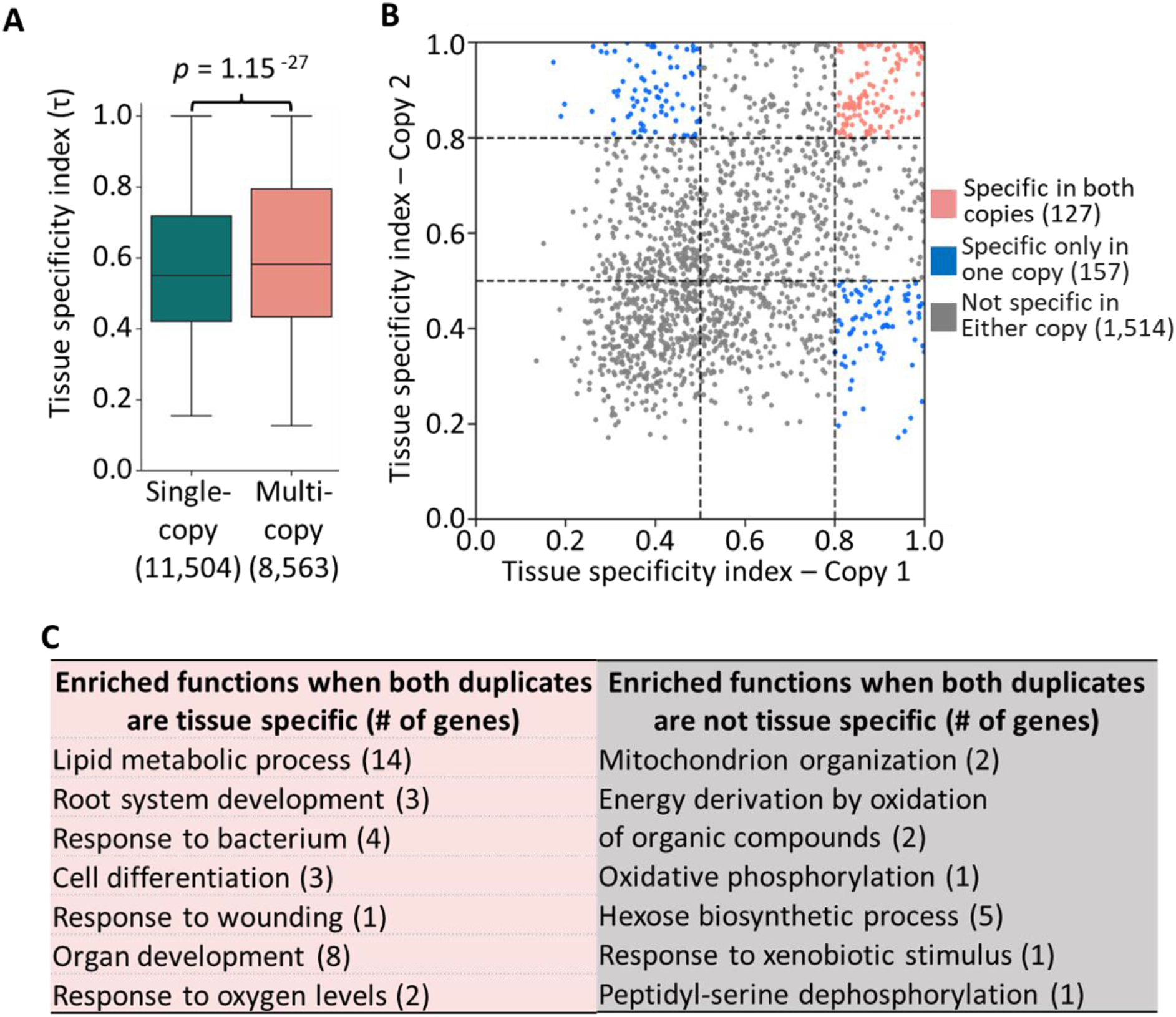
Duplicated genes show higher average tissue specificity compared to single-copy genes. **[A]** Expression specificity measured by tissue specificity index (τ) for single- and multi-copy genes *p*-value based on Wilcoxon test applied to two groups. **[B]** Preferential expression between two copies in a duplicated gene among all samples. Dash lines represent τ ≥ 0.5 and 0.8 for each copy. **[C]** Functional processes enriched among duplicated genes where both copies are tissue specifically expressed (left) or neither copy is tissue specifically expressed (right).

### Spatiotemporal expression of differentially expressed genes in response to high salinity

To investigate how differentially expressed genes (DEGs) in response to salt are coordinated across tissues and developmental stages in *S. parvula*, we identified DEGs at a *p*_*adj*_ ≤ 0.01 using DESeq2 (Table S4). Additionally, we included salt treated transcriptome data from a previously published dataset to expand the number of salt treatment conditions tested in the current study. This included 150 and 250 mM NaCl given to 4-week old bulk root and shoot tissue harvested at 0, 3, and 24 hours following treatment (Tran *et al*., 2022). Our analysis includes 27 different salt treatments given from 3 hours to three weeks, two different NaCl concentrations, and single NaCl and multiple-salt combinations compared to their respective control conditions examined at different developmental stages and tissues (Figure 8). The magnitude of the transcriptomic response (assessed as a fold change distribution detected for DEGs) per sample was highest for samples treated with multiple salts compared to single NaCl treatments (upper plot; Figure 8A). *S. parvula* did not seem to respond drastically to an increase in the salt concentration from 150 to 250 mM or 150 mM NaCl treatment duration increased from three hours to three weeks. However, within a single NaCl concentration, seedlings responded at a higher magnitude more than mature tissues (upper plot; Figure 8A). When assessing the number of unique DEGs expressed per sample under salt compared to its control condition flowers, siliques, and seeds had the highest number of unique DEGs per sample. Notably, multiple-salt treated samples that showed the highest response had comparable number of unique DEGs compared to other samples (lower plot; Figure 8A).

**Figure 8.**
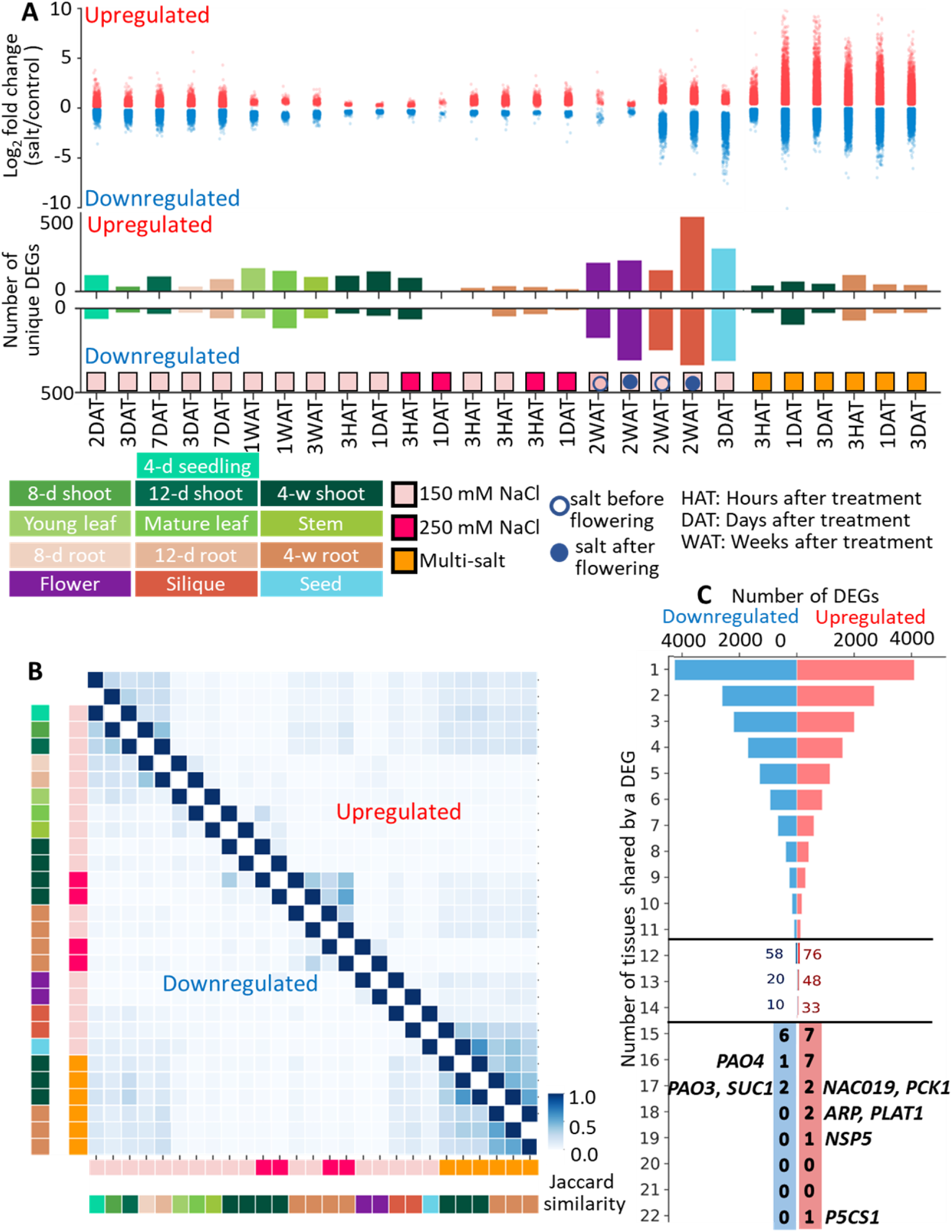
Spatiotemporal expression in response to salt. **[A]** Upper plot: magnitude of salt responses in fold changes from differentially expressed genes (DEGs). Lower plot: number of DEGs per sample compared to its control condition. **[B]** Transcriptome responses to salt measured as Jaccard similarity among all samples. Colored bars on x- and y- axes indicate the sample types and treatments. **[C]** The number of DEGs in response to salt found as a shared DEG among samples. The y-axes represents three parts to highlight the number and type of genes with fewer genes shared by increasing number of samples.

We assessed the shared salt response among samples using Jaccard similarity calculated for DEGs identified for all pairs of salt treated samples (Figure 8B). This analysis highlighted that there were no two salt transcriptomes highly similar to one another with every sample consisting of a high number of distinct DEGs. Despite the overall uniqueness of the salt responsive transcriptomes, the samples treated with multiple salts and young seedlings had more similarities within their group than similarities within any other tissue group, developmental stage, or salt treatment class (Figure 8B). There were 4,101 upregulated genes and 4,261 downregulated genes found to be a DEG in just one salt treated sample compared to its control representing the most uniquely expressed set of DEGs. Less than 5% of DEGs were found in more than half of the samples and none of them were shared by all samples. The most commonly expressed DEG found in 22 salt treated samples out of 27 was *pyrroline-5-carboxylate synthetase 1* (*P5CS1*) which is a highly induced gene during salt stress in plants. (Shinde *et al*., 2016) (Figure 8C).

Given the high dissimilarity between salt responsive transcriptomes, we sought to test if those transcriptomes shared any stress responsive pathways or functional groups distributed across tissues and developmental stages. Figure 9 summarizes the functional groups represented by upregulated DEGs in our study. Figure S6A provides the equivalent summary for downregulated DEGs. Details about these enriched functional groups are given in Table S5. These highlighted that there was a significant convergence in the functional responses from the majority of tissues and developmental stages that responded to salt. We did not detect an over-representation for stress responsive functional groups in any of the tissues and developmental stages (Figure S6B). Notably, 20%-30% of DEGs from each tissue were not annotated by any GO functions likely indicating a gap in the functional associations recognized for genes involved in salt stress responses in extremophytes.

**Figure 9.**
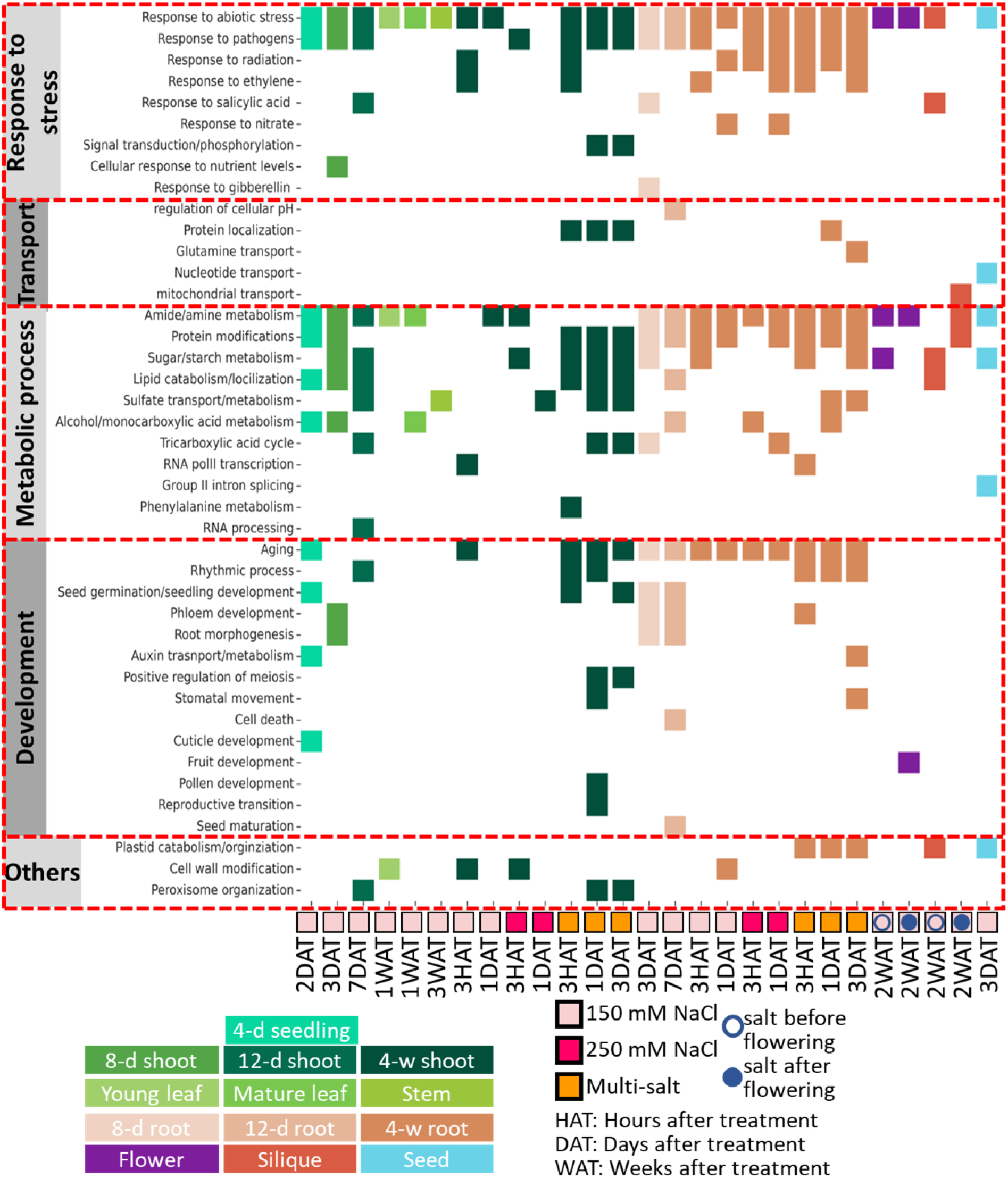
Functional processes enriched among upregulated differentially expressed genes in different samples. Enriched GO terms are grouped into and labelled with different functional categories.

### Transcriptome network reprograming in the presence of salt

To investigate the salt stress-induced changes in gene co-expression, we compared gene co-expression networks constructed separately from control and treated samples by calculating the ratio of edges for each gene between the two conditions. We observed a dramatic decrease in network edges derived from treated samples compared to that from control samples for most genes (Figure S7A), suggesting that salt stress reduces the number of co-expressed genes at the global level. As transcription factors play a major role in defining the co-expression landscape, we next searched for transcription factors with altered co-expression connectivity in response to salt stress. This led to the identification of 816 transcription factors with losing co-expression connections and 116 gaining connections upon salt stress (Figure S7B). Distinct functional groups were found to be represented by these transcription factors with stress response-related functions and development-related processes enriched among the ones with increased and reduced connections, respectively (Figure S7C). To further investigate salt stress-induced changes in co-expressed genes of these transcription factors, we categorized the 116 transcription factors with gained connections mapped to their preferential expression patterns in tissues and developmental stages and examined their co-expression neighborhood (Figure 10).

**Figure 10.**
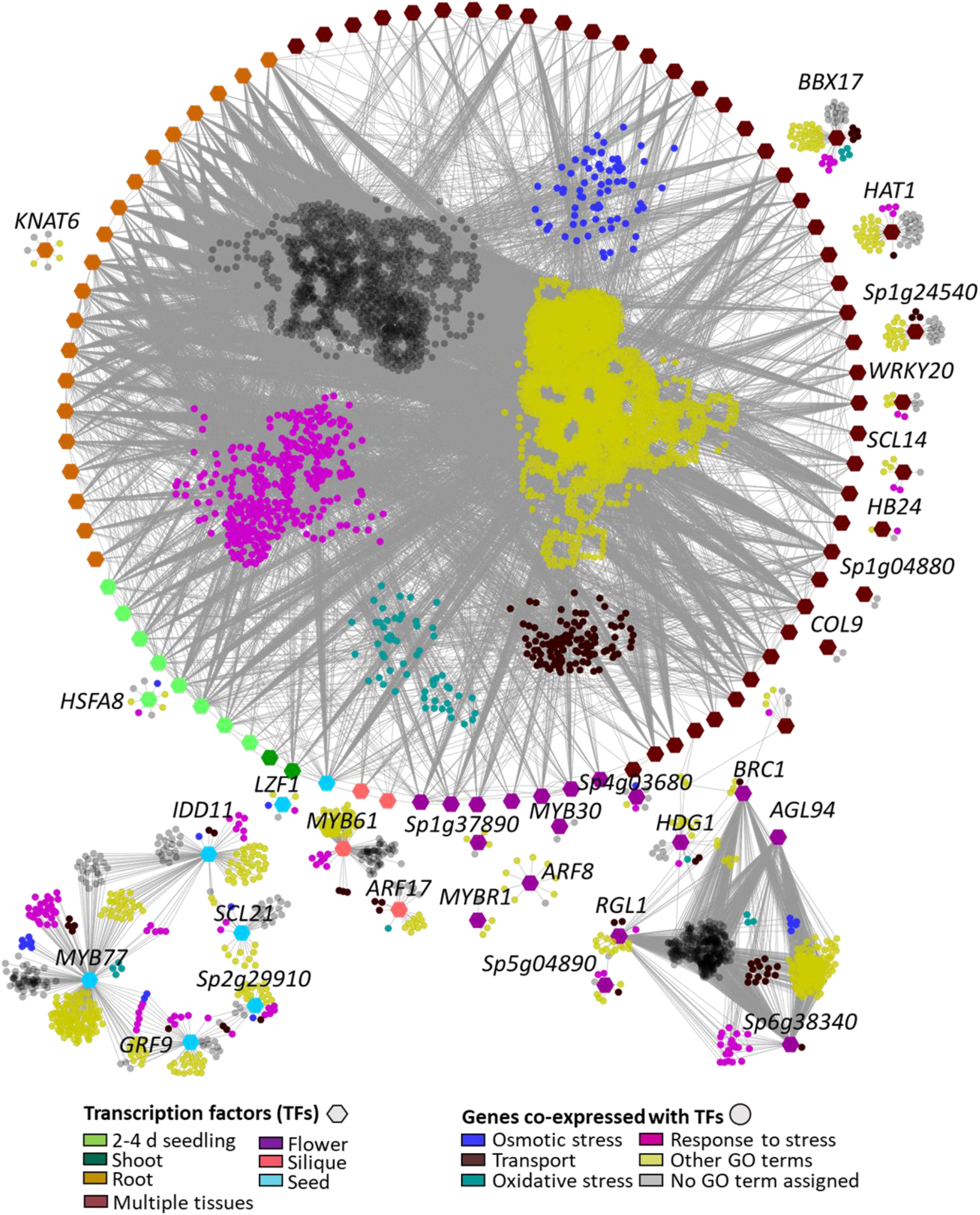
Co-expression network for transcription factors with increased connections in response to salt. Nodes indicate either transcription factors as hexagons or their co-expressed genes as circles. Edges represent significant correlations between the transcription factors and their co-expressed genes. Transcription factors and their co-expressed genes are colored based on tissues they are preferentially expressed in and the functional annotations, respectively.

Co-expression networks that were associated with transcription factors preferentially expressed in reproductive tissues, such as flowers and seeds, barely showed any overlaps with these from other tissues, indicating distinct salt responses in these reproductive tissues. Regardless of the tissue specificity of the co-expression networks, stress response-related genes appear to constitute a significant proportion of networks. It is also worth noting that the majority of the co-expressed genes tend to be associated with functions not related to stress responses, which can provide valuable insights for processes that accompany the stress responses in *S. parvula* to maintain its growth under salt stress considering that *S. parvula* is barely affected by the given salt treatments (Figure 10).

## DISCUSSION

This study adds a novel transcriptome resource for the extremophyte model *S. parvula* and provides insights into how spatiotemporal gene expression profiles are coordinated across tissues and developmental stages when responding to high salinity. Based on transcriptomes generated for bulk shoots and roots from 4-week-old plants, previous studies had shown that *S. parvula* was preadapted in its transcriptomic responses to salinity changes and showed fewer but highly coordinated responses under salt compared to *A. thaliana* (Oh *et al*., 2014; Pantha *et al*., 2021; Tran *et al*., 2022). Recent physiological assessments across tissues and developmental stages in *S. parvula* implied that we may have missed salt responsive gene networks that are not detected at the mature vegetative growth stages or with short-term salt treatments (Tran *et al*., 2021). We have now significantly expanded the past resources to include 115 transcriptomes sequenced from 35 tissue and developmental stages examining their responses before and after 27 salt treatments in our current study.

*A. thaliana* has been a quintessential genetic model primarily because of the collective resources contributing to its spatiotemporally resolved gene expression data and follow up functional studies done to confirm new gene functions (Krishnakumar *et al*., 2015). Our study leverages the phylogenetic proximity of *A. thaliana* to *S. parvula* when attributing functional significance to tissue or developmental stage-preferentially expressed, differentially expressed, or coordinately co-expressed in networks in response to high salinity. Thereby, we were able to identify highly studied as well as functionally still uncharacterized genes categorized into pathways, functional groups, and networks spatiotemporally distinct in their expression in an extremophyte model when experiencing salinity levels toxic to most crops (Flowers and Colmer, 2015).

Reproductive tissues had the lowest number of expressed genes compared to other tissue types, while seedlings showed the highest magnitude of responses to salt stress among other developmental stages. Yet, seedlings had fewer uniquely expressed genes and seeds had the highest tissue specifically expressed genes compared to all other samples. This highlights the importance of examining both the differential expression and tissue specificity when deducing stress responsive networks in *S. parvula* (Figure 2, 4, and 8).

Stress experiments targeting one tissue type (often roots) or a single developmental stage (often seedlings) have immensely contributed to our current mechanistic understanding about genetic responses to salt stress in plants (Brady *et al*., 2007; Song *et al*., 2016). However, such studies cannot be readily extrapolated to identify stress responsive networks coordinated across tissues and developmental stages to result in the successful completion of a plant lifecycle when experiencing environmental stresses. A transcriptome atlas aids in bridging the gaps between a functional gene group highly characterized at a specific condition and its members diversified and connected with new associations at a global level. Our results support the view that salt tolerance is spatiotemporally highly coordinated (Geng *et al*., 2013) by illustrating that individual transcriptome responses had less than 5% of DEGs shared among half of the salt treated samples we examined (Figure 8).

A typical transcriptomic study will be able to detect differentially expressed genes in response to stress albeit the uncertainty of their prevalent expression in other tissues or different devolvement stages. However, the expression specificity of a gene or its changes in specificity in response to stress cannot be deduced with a few transcriptome profiles. A transcriptome atlas aids to close this gap and allows the identification of marker genes at variable specificity. We were able to create a reference gene list with high specificity in their expression for diverse tissues and developmental stages for *S. parvula* (Figure 4 and 10). This can serve as a valuable resource when deciding suitable gene modifications with minimal off-target effects for salt-resilient growth (Zhao *et al*., 2020; Rawat *et al*., 2022).

In summary, we have generated a transcriptome atlas for *S. parvula* which could serve as a critical resource to explore innovative stress responsive networks evolved in an extremophyte. Our analysis revealed that *S. parvula* shows unique salt responsive gene networks that are highly coordinated across tissues and developmental stages but connected with shared functions associated with environmental stresses.

## EXPERIMENTAL PROCEDURES

### Plant growth and sampling

*Schrenkiella parvula* (ecotype Lake Tuz, Turkey; Arabidopsis Biological Resource 575 Center/ABRC germplasm CS22663) seeds were surface sterilized, stratified, and grown on plates or hydroponically as described in (Wang *et al*., 2021). Plants in soil were grown as described in (Wang *et al*., 2019). See Table S1 for plant growth, treatment, and sampling details for RNAseq libraries.

### Transcriptome profiling and data preprocessing

Total RNA was extracted using QIAGEN RNeasy Plant Mini Kit (QIAGEN, Hilden, Germany) from all samples except for seeds, with an on-column DNase treatment to remove contaminating DNA. To isolate total RNA from seed samples, we first followed an optimized method for seeds (Kanai *et al*., 2017) to remove compounds such as oils, proteins, and polyphenols, and subsequently performed RNA extraction using QIAGEN RNeasy Plant Mini Kit. About 1 μg of total RNA per tissue type at a quality of RNA integrity number ≥ 6 based on a Agilent 2100 Bioanalyzer (Agilent Technologies, CA, USA) were used to generate RNA-Seq libraries. For each tissue type, developmental stage and treatment, at least three biological replicates were used for RNA isolation. RNAseq libraries were constructed with True-Seq stranded RNAseq Sample Prep Kits (Illumina, San Diego, CA, USA), multiplexed, and sequenced on a Novo-Seq platform (Illumina) at Joint Genomics Institute (JGI) at Berkeley, CA, USA. A minimum of 50 million reads with 151-nucleotide paired-reads per sample were generated for a total of 107 RNA-seq samples.

Quality testing and initial data preprocessing were conducted at JGI, Berkeley, CA, USA. During this process, BBDuk (v38.94) was used to remove contaminants, trim reads that contained adapter sequence, remove homopolymers of G’s of size five or more at the ends of the reads and to trim the right end of the read when base quality is below Phred quality of 6. An additional filtering step was performed to remove reads if the read contains more than one N base, or if the average quality score across the read was less than 10, or if read length was ⩽49. The cleaned data were used for all downstream analyses.

### Read mapping and quantification

RNAseq reads after quality filtering from each sample were mapped to *S. parvula* v2.2 transcriptome using Salmon (v1.3.0) (Patro *et al*., 2017) with the -gcBias option to normalize for local GC content and quantify transcript abundance. Library type was automatically inferred by setting the “-l” parameter as “A” option. Normalized expression as transcript per million (TPM) values obtained from Salmon were used to filter lowly expressed genes and highly variable genes across replicates using custom python scripts based on the following criteria: 1) the expression of a gene was greater ≥ 2 TPM in at least one tissue; 2) the z-score of the expression of a gene between replicates was ≥ 2 in at least 2/3 of the replicates. The filtered gene list was used for all downstream analyses except for differential expression quantified with DESeq2.

### Tissue specificity index calculation

Tissue specificity of gene expression was measured using Tissue Specificity Index (τ) for all genes following the approach described in (Yanai *et al*., 2005). For a given gene, τ ranges from 0 to 1 with higher values representing higher tissue specificity. τ were computed separately for control and treated samples. Four-week-old shoot and root samples were excluded from this analysis. Tissue specific markers were identified by filtering genes with a τ ≥ 0.8 among the control tissues. Salt stress-induced changes in τ were evaluated by calculating the difference in τ between control and salt treated samples for each gene. For this study, we considered a gene losing tissue specificity in response to salt stress if Δ τ _(control - treated)_ ≥ 0.25 or gaining tissue specificity if Δ τ _(treated - control)_ were ≥ 0.25.

### Differential expression analysis

DESeq2 (v1.34.0) (Anders and Huber, 2010) was used to normalize for library size and identify differentially expressed genes across different tissues. We used Taximport (v1.24.0) (Soneson *et al*., 2015) in R to convert TPM values derived from Salmon to raw count data as the input for DESeq2. A gene was considered to be differentially expressed between treatment groups with an adjusted *p*-value ≤ 0.01. The overlaps in differentially expressed genes (DEGs) between different tissues were calculated as Jaccard similarity index for all tissue pairs. The Jaccard index between any two tissues will be 1 if they have identical DEG sets, but 0 if they do not share any DEGs.

### Gene co-expression module identification and network construction

Gene co-expression modules among control samples, except the multiple salt treated shoot and root samples, were identified using weighted gene co-expression network analysis (WGCNA) (v1.69) (Langfelder and Horvath, 2008) with “mergeCutHeight” set to 0.3 and “scale” to 9. To enable detailed comparisons of network properties between treated and control samples, gene co-expression networks were constructed using normalized TPM values independent of WGCNA. We first calculated the Pearson correlation of expression for all possible gene pairs using Hmisc (v4.7-0) on R and then identified significant correlations with an adjusted *p*-value ≤ 0.05, which were considered as signed correlations. We considered any gene pairs with a signed correlation ≥ 0.95 as a co-expressed pair. Co-expression networks were built separately for control and treated samples and the number of edges for each node was calculated for both networks. Transcription factors with altered connections were extracted if the log_2_ ratio of connections associated with the transcription factor between the treated and control conditions was larger than 1 or smaller than -1. To further determine the tissue attributes of these transcription factors, we categorized all tissues into broader tissue groups as follows: seedling group included 2-day-old and 4-day-old seedlings; shoot tissue group included shoots from 8-day-old and 12-day-old seedlings, stems, and leaves; root tissue group included roots from 8-day-old and 12-day-old seedlings. All reproductive tissues, including flowers, seeds, and siliques, remained as independent groups. For each transcription factor, we calculated the fraction of its expression in each tissue group over its total expression from all tissue groups and determined the tissue groups that independently contributed to more than 25% of the total expression, which were considered as the tissue groups the given transcription factor was preferentially expressed in. Co-expression networks were visualized using NetworkX (v2.8.1) and Cytoscape (v3.7.2).

### Gene ontology annotation and analysis

BiNGO (v3.0.5) (Maere et al., 2005) was employed to identify enriched functional terms among any gene sets of interest using their Arabidopsis orthologs. We further reduced the number of non-informative GO terms and the redundancy between enriched GO terms using GOMCL (v0.0.1) (Wang *et al*., 2020) followed by manual curation of broader categories based on their functional relevance. To assign individual *S. parvula* genes to GO categories, we first downloaded GO annotation for Arabidopsis from The Arabidopsis Information Resource (TAIR) (https://www.arabidopsis.org/download_files/GO_and_PO_Annotations/Gene_Ontology_Annotations/) (Berardini *et al*., 2004) and transfer GO annotations from Arabidopsis to *S. parvula* using orthologous gene pairs reported in (Oh and Dassanayake, 2019). Genes annotated with GO categories “transmembrane transport” (GO:0055085), “response to oxidative stress” (GO:0006979), “response to osmotic stress” (GO:0006970) were extracted separately. If a gene was annotated with multiple above-mentioned categories, or with the GO term “response to stress” (GO:0006950), we assigned it to the “response to stress” group. Rest of the genes that cannot be annotated with any of the GO terms above were divided into two categories, either with at least one other GO annotation or without any GO annotation.

### Duplicated genes and transcription factor identification

Duplicated genes in *S. parvula* were identified by first obtaining ortholog groups between five Brassicaceae species, including *S. parvula*, as OrthoNets from a previous study (Oh and Dassanayake, 2019) and searching for OrthoNets with no more than twenty members. We consider a *S. parvula* gene as a multi-copy gene if the OrthoNet where the given gene was found contains at least one more *S. parvula* gene while there is only one gene from each of the rest of Brassicaceae species. Potential transcription factors in *S. parvula* were downloaded from Plant Transcription Factor Database (http://planttfdb.gao-lab.org/).

## DATA AVAILABILITY STATEMENT

All relevant data can be found within the manuscript and its supporting materials. The RNAseq Illumina reads for all samples used to create the transcriptome atlas is available at the U.S. Department of Energy Joint Genome Institute Genome portal accessible at https://genome.jgi.doe.gov/portal/SchparEProfiling/SchparEProfiling.info.html under Project ID: 1263770 and Project Title: A transcriptome atlas enabling discovery of genes evolved as adaptations to environmental stress in a model extremophyte, *Schrenkiella parvula*. The RNAseq Illumina reads used for bulk root and shoot samples treated with 150 mM NaCl from four-week old plants have been deposited in the Sequence Read Archive at the National Center for Biotechnology Information (http://www.ncbi.nlm.nih.gov/sra) under accession number PRJNA63667. The spatio-temporal gene expression browser developed in this study for *Schrenkiella parvula* is available at https://www.lsugenomics.org/resources.

## ACKNOWLEDGEMENTS

RNA sequencing for this study was supported by Department of Energy Joint Genome Institute (JGI) Award DE-AC02-05CH11231. Authors thank Dr. Kerrie Barrie and other staff members at JGI who coordinated sample processing and sequencing with constant communication between labs, Ms. Prava Adhikari for assistance in RNA extractions, and Mr. Richard Garcia for providing helpful comments. This work was additionally supported by the US National Science Foundation awards MCB-1616827 and IOS-EDGE-1923589, and US Department of Energy BER-DE-SC0020358 awarded to MD. CW, GW, PP, and KT were also supported by an Economic Development Assistantship award from Louisiana State University. We acknowledge LSU High Performance Computing services for providing computational resources for this study.

## AUTHOR CONTRIBUTIONS

CW, GW, PP, and KT conducted wet lab experiments from plant growth to extracting RNA. CW performed bioinformatics analyses. MD developed the experimental design and supervised the overall project. CW, GW, PP, KT, and MD interpreted results and wrote the article.

## SUPPORTING INFORMATION

**Figure S1.**
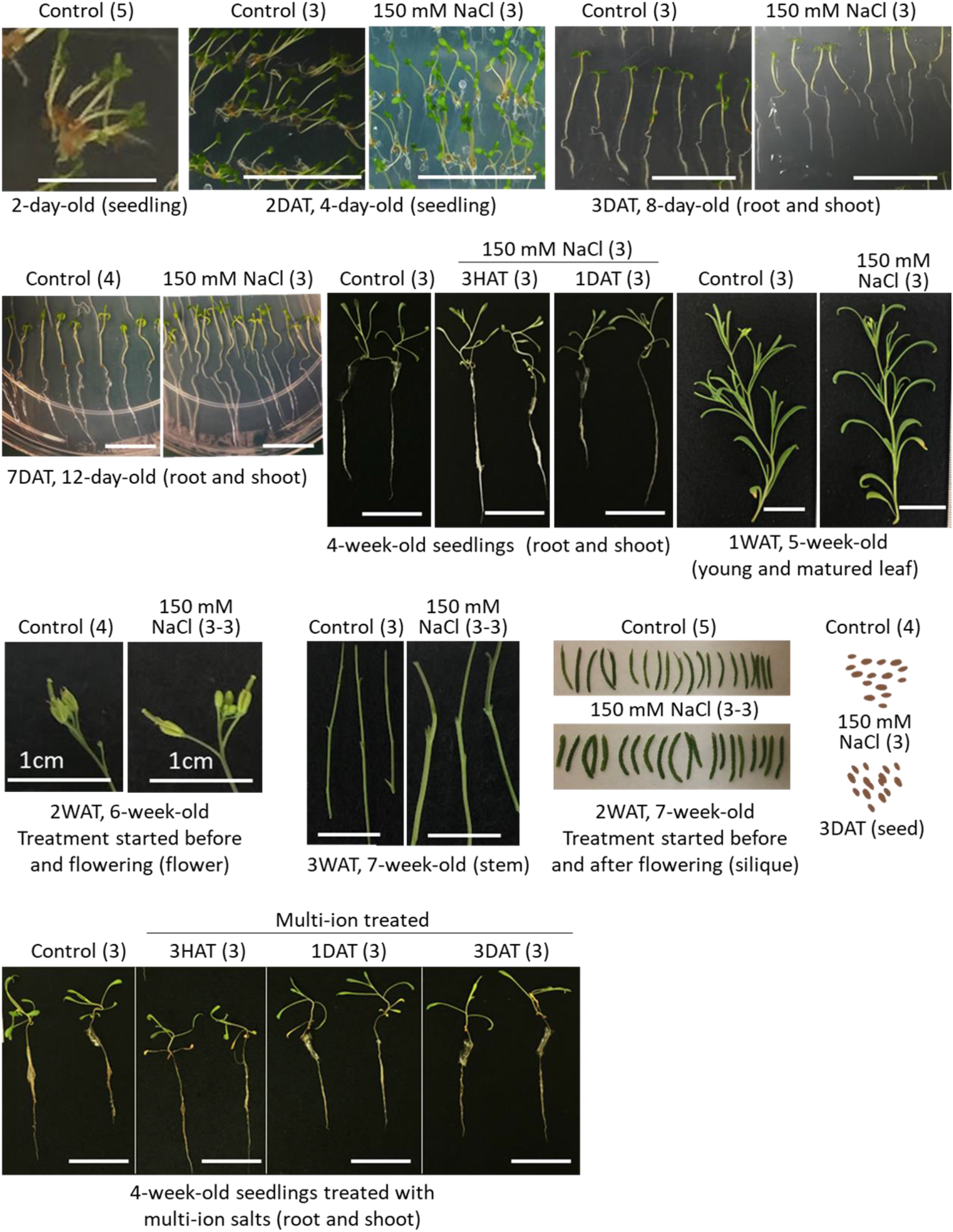
Images of *Schrenkiella parvula* tissues/organs used to generate the transcriptome atlas

**Figure S2.**
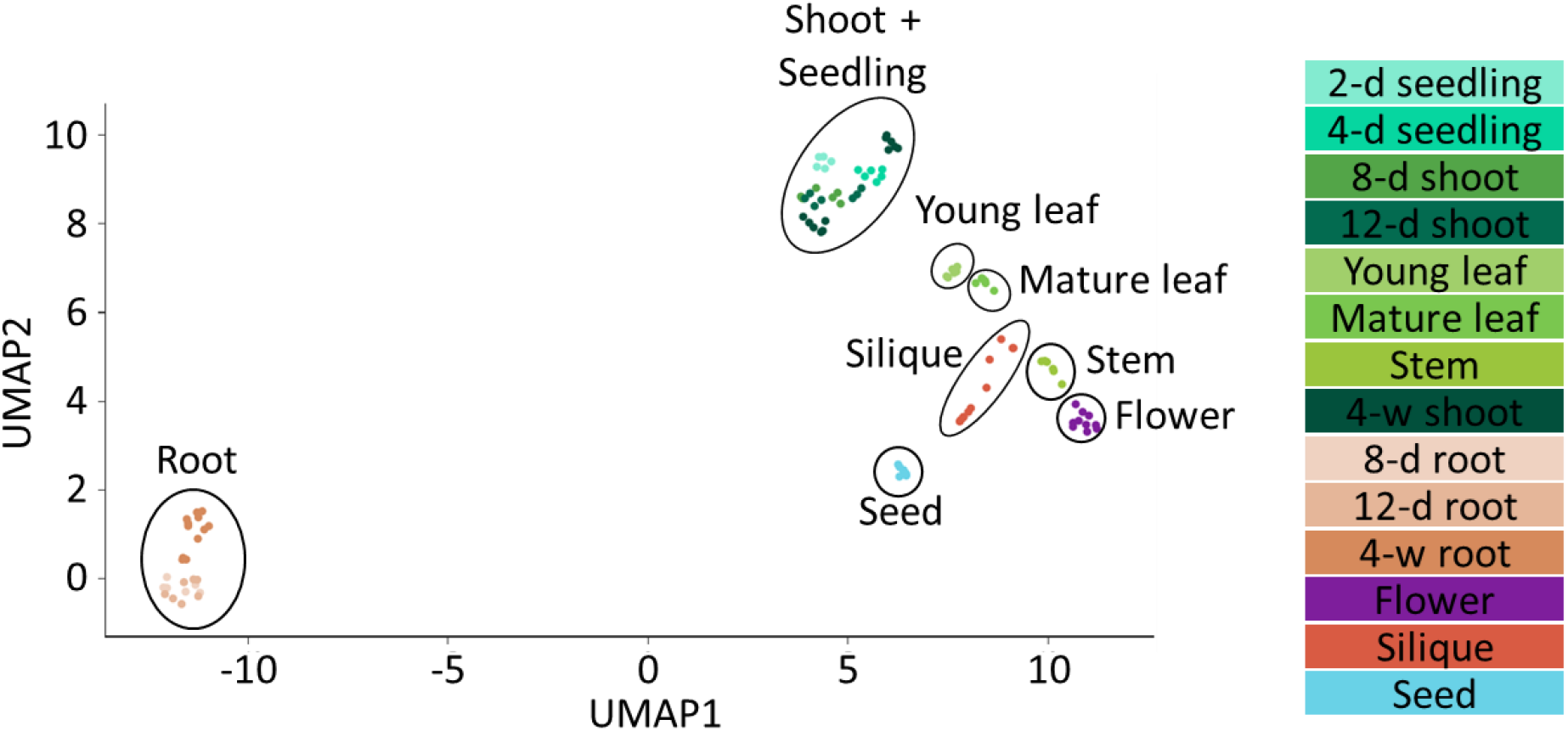
UMAP visualization of *S. parvula* transcriptomes

**Figure S3.**
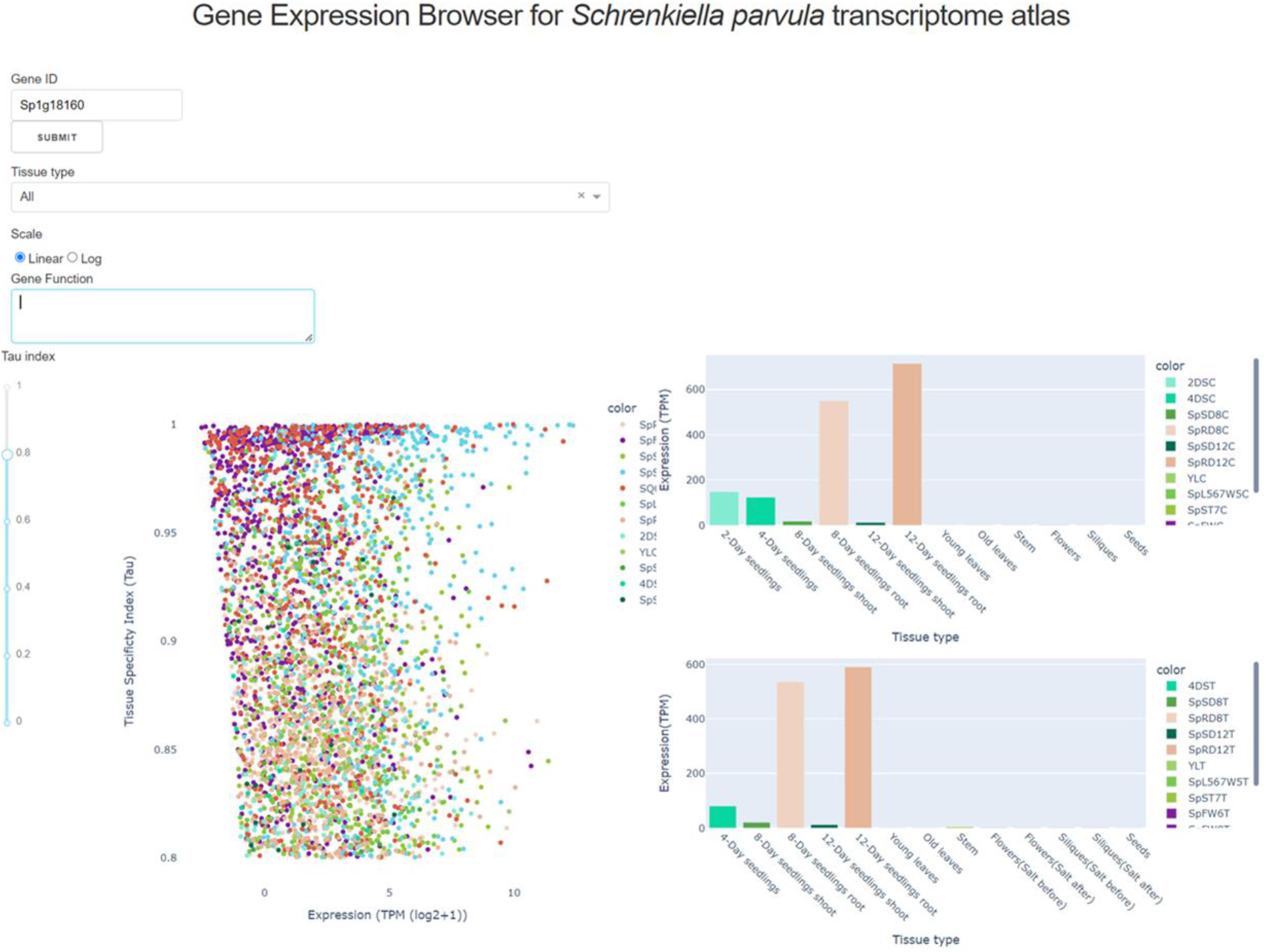
Screenshot of *S. parvula* transcriptome atlas browser

**Figure S4.**
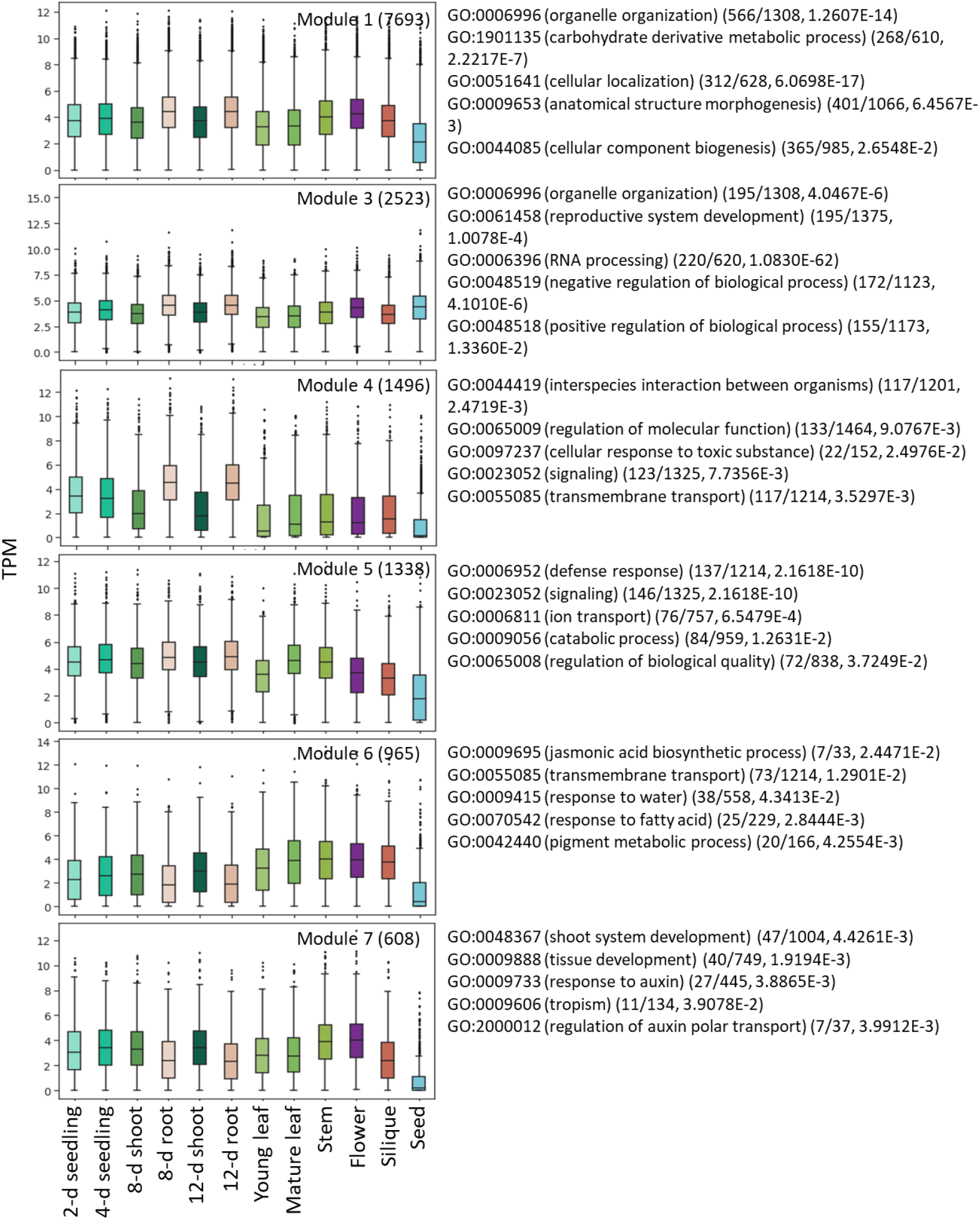
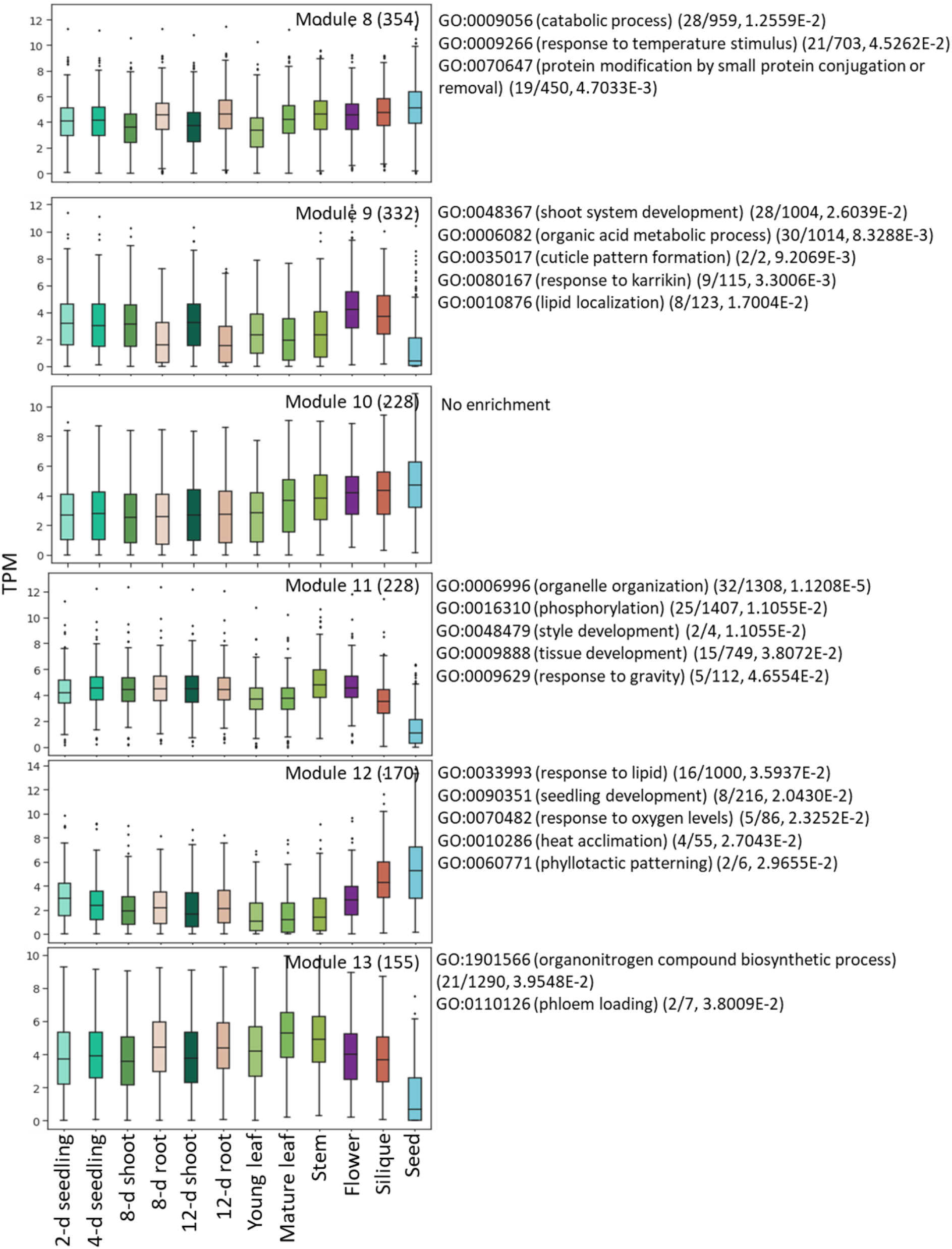
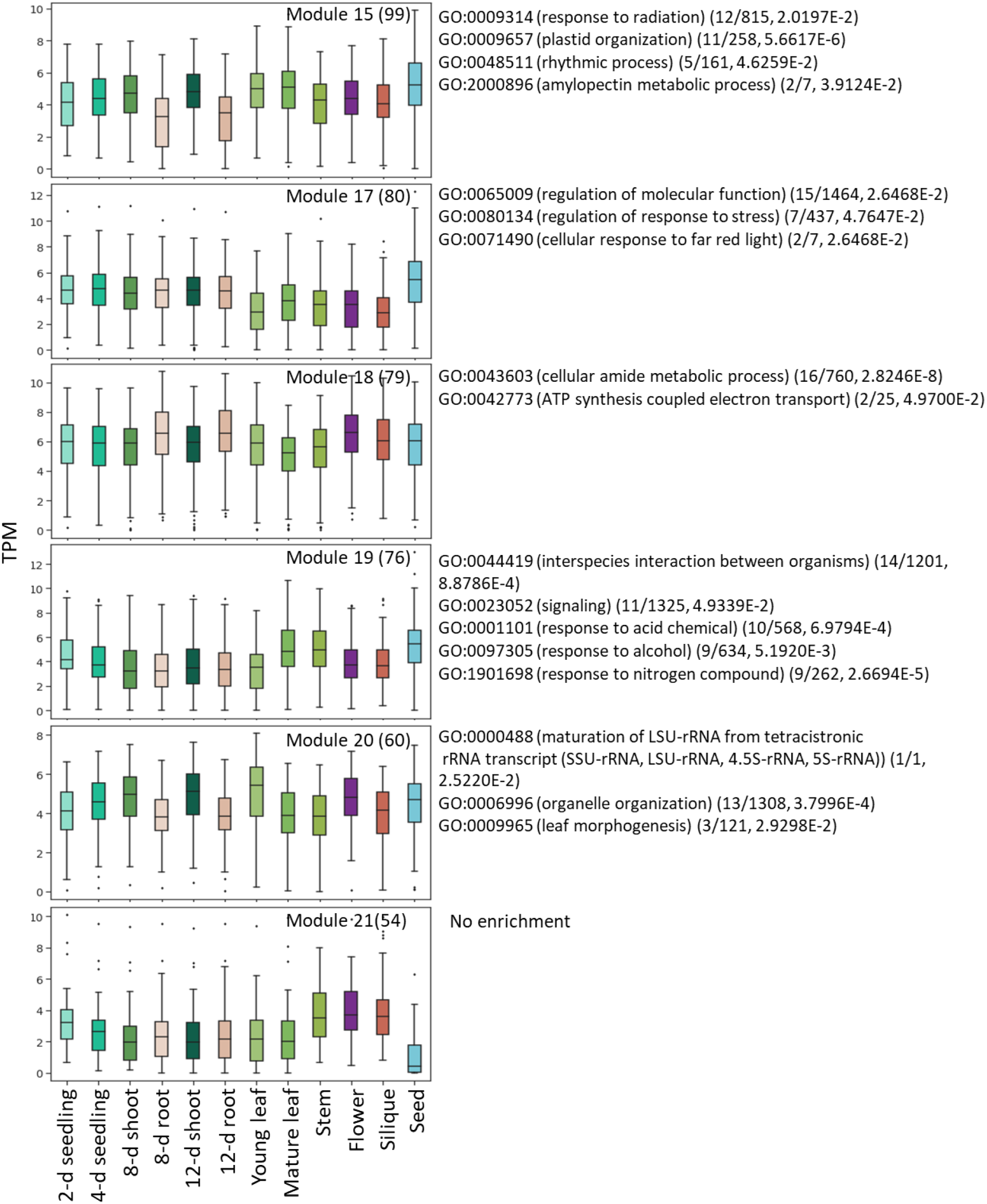
Expression distribution of co-expression modules across tissues and developmental stages and functional processes enriched in these modules

**Figure S5.**
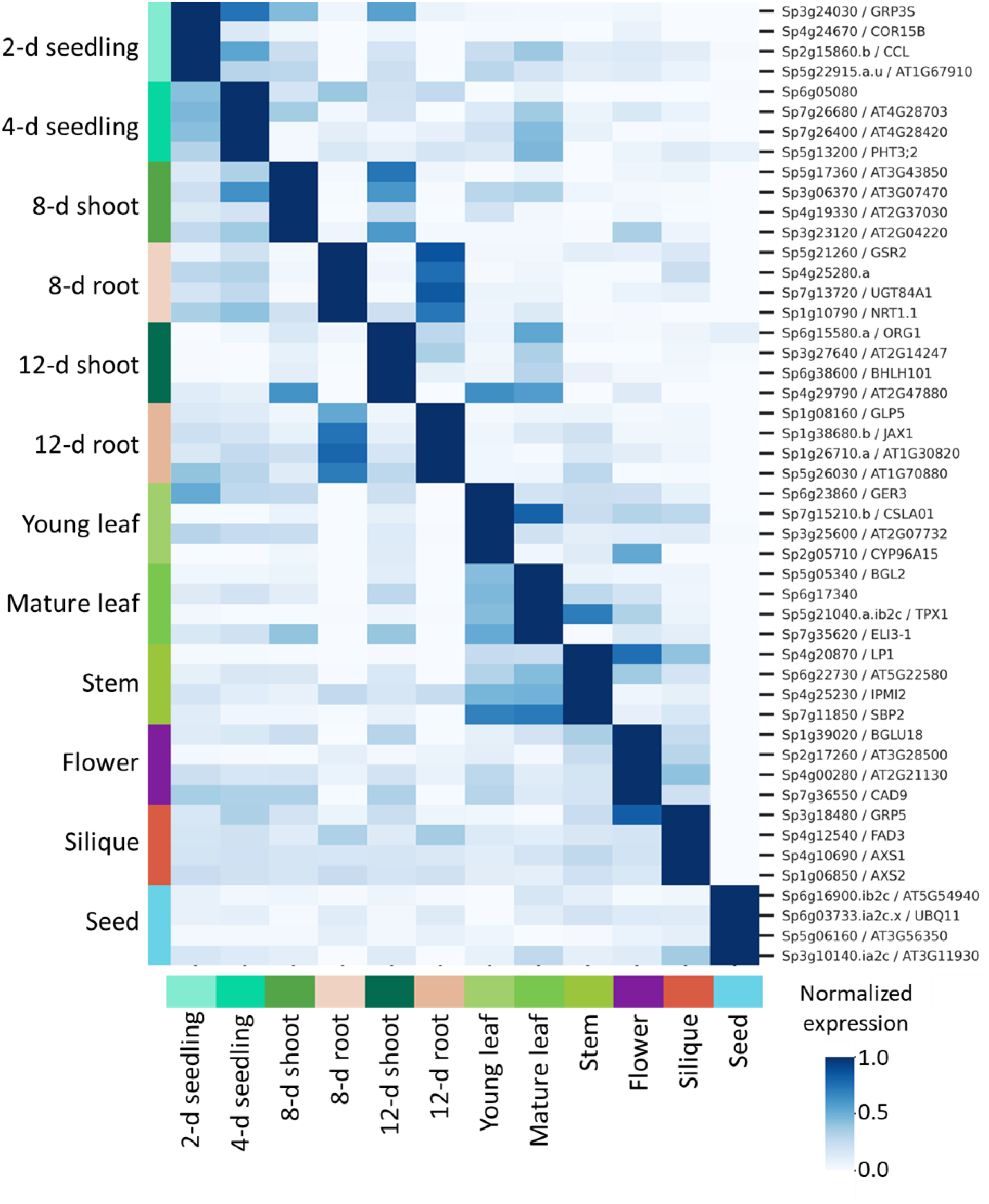
Normalized expression of top four marker genes identified for each tissue and developmental stage

**Figure S6.**
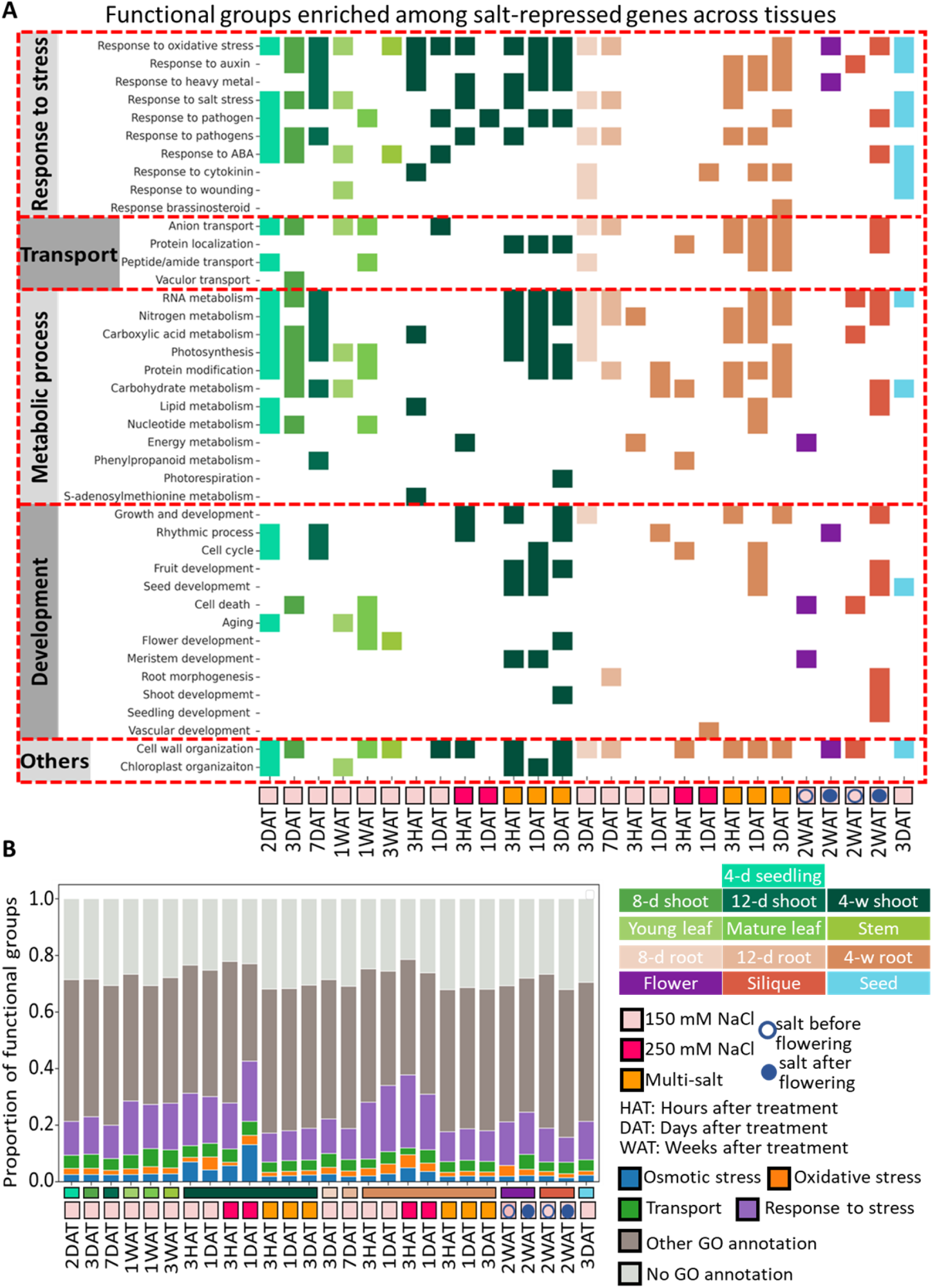
Functional processes represented by DEGs in different samples

**Figure S7.**
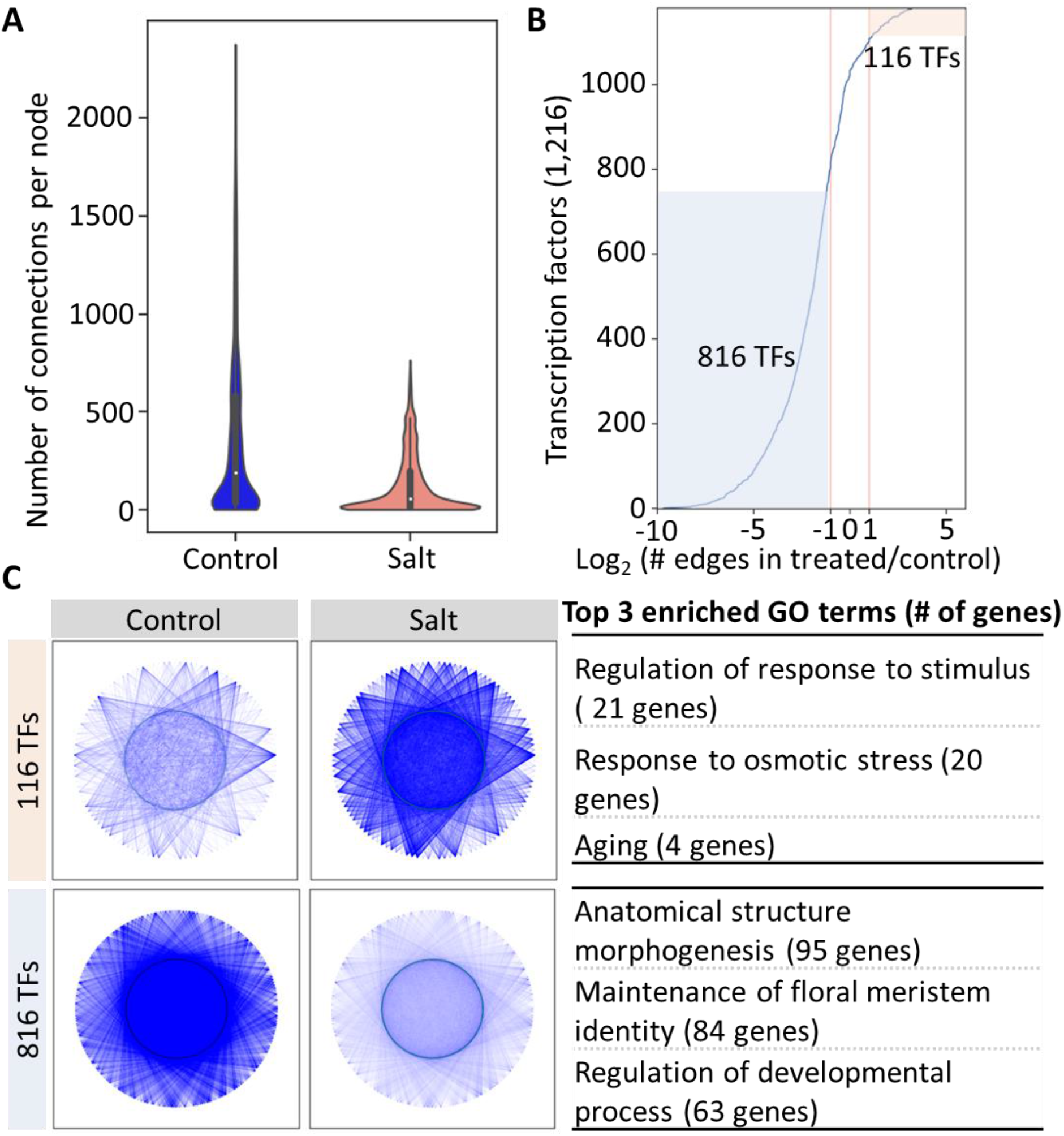
Effect of salt stress on gene co-expression networks

Table S1. Sample metadata

Table S2. Normalized gene expression

Table S3. Tissue specificity index values

Table S4. Differential expression analysis results

Table S5. Functional processes enriched in DEGs

